# Metabolic Quadrivalency in RSeT Human Embryonic Stem Cells

**DOI:** 10.1101/2024.02.21.581486

**Authors:** Kevin G. Chen, Kyeyoon Park, Dragan Maric, Kory R. Johnson, Pamela G. Robey, Barbara S. Mallon

**Author notes:** **Correspondence:** (K.G.C.), (B.S.M.).

## Abstract

One of the most important properties of human embryonic stem cells (hESCs) is related to their pluripotent states. In our recent study, we identified a previously unrecognized pluripotent state induced by RSeT medium. This state makes primed hESCs resistant to conversion to naïve pluripotent state. In this study, we have further characterized the metabolic features in these RSeT hESCs, including metabolic gene expression, metabolomic analysis, and various functional assays. The commonly reported metabolic modes include glycolysis or both glycolysis and oxidative phosphorylation (i.e., metabolic bivalency) in pluripotent stem cells. However, besides the presence of metabolic bivalency, RSeT hESCs exhibited a unique metabolome with additional fatty acid oxidation and imbalanced nucleotide metabolism. This metabolic quadrivalency is linked to hESC growth independent of oxygen tension and restricted capacity for naïve reprogramming in these cells. Thus, this study provides new insights into pluripotent state transitions and metabolic stress-associated hPSC growth *in vitro*.

## INTRODUCTION

In the past two decades, various pluripotent states have been identified in human pluripotent stem cells (hPSCs) both *in vitro* and *in vivo*, including naïve, formative, and primed pluripotency [reviewed in (Dong et al., 2019; Smith, 2017; Weinberger et al., 2016; Zhou et al., 2023; Zimmerlin et al., 2017)]. Naïve and primed pluripotent stem cells represent the states of pre-and post-implantation epiblasts, respectively. These two states can be interconverted in a laboratory setting. Formative pluripotency is a state between the naïve and primed states of epiblast, which has a decisive role in determining pluripotency transitions and multi-lineage competence (Smith, 2017). The establishment of various human pluripotent states has the potential to greatly impact our knowledge of early human embryonic development, enhancing the growth of hPSCs in vitro, replicating disease models, discovering novel drugs, and advancing regenerative medicine (Chen et al., 2018; Chen et al., 2021; Cherry and Daley, 2013; Zhou et al., 2023).

Despite the existence of various protocols to generate hPSCs with different pluripotent states, there are increasingly intense efforts to derive naïve human pluripotent stem cells based on the commercially available RSeT medium (www.stemcell.com) (Collier et al., 2017; Kilens et al., 2018; Liu et al., 2017; Szczerbinska et al., 2019). Nevertheless, the characteristics of RSeT cells and their pluripotent states have not been thoroughly studied under various growth conditions. In two recent studies, we reported detailed characterization of RSeT hESCs and defined a unique pluripotent state downstream of formative pluripotency, similar to primed hPSCs (Chen et al., 2024; Johnson et al., 2021). We reported that the RSeT hESCs showed multiple novel characteristics, such as autonomous growth in the absence of hypoxia, heterogeneous expression of primed and naïve markers, and varied reliance on FGF2, JAK, and TGFβ signaling in a cell-line-specific manner ( Chen et al., 2024). These standout features differentiate these cells from those of primed and naïve hPSCs, which highlights the need for in-depth investigation on other requirements of hPSCs.

It has been shown that metabolic requirements are an important feature for pluripotent stem cell growth, state transitions, and self-renewal (Takashima et al., 2014; Zhou et al., 2012). The metabolic needs of hPSCs encompass the use of anabolic precursors to facilitate the duplication of their genetic material and cellular components, as well as the consumption of ATP for energy (Tsogtbaatar et al., 2020). From this, this is evident that primed cells predominantly utilize glycolysis, and naive cells employ the bivalent mode [(including glycolysis and oxidative phosphorylation (OXP)] (Somasundaram et al., 2020; Takashima et al., 2014; Tsogtbaatar et al., 2020; Wu and Izpisua Belmonte, 2015; Zhang et al., 2018; Zhou et al., 2012). Furthermore, hPSC-derived metabolites can directly and indirectly impact epigenetic and transcriptional landscapes, ultimately influencing self-renewal properties (Somasundaram et al., 2020). Therefore, gaining insight into the roles of metabolic pathways offers a basis for understanding the factors that control the growth potential and the governance of the unique pluripotent states in these RSeT cells.

In this study, we have further elucidated the metabolic attributes of RSeT hESCs by deciphering metabolic gene expression, metabolomic analysis, and by performing several functional tests. We focus on mapping the diverse metabolites in RSeT hESCs in order to thoroughly evaluate their metabolic states. Moreover, we seek to determine the relationship between the metabolic behavior of RSeT hESCs and their limited capacity for metabolic reprogramming, as well as their ability to sustain the hypoxia-independent growth in vitro. We demonstrate that RSeT hESCs exhibit a striking metabolic quadrivalency, which includes the previously mentioned metabolic bivalency, in the presence of fatty acid oxidation (FAO) and notably altered or imbalanced nucleotide metabolism. This new metabolic mode seems to be linked to acquiring their unique pluripotent state and growth potential.

## RESULTS

### Changes in metabolic gene expression patterns in RSeT hESCs

RSeT medium is based on naive human stem cell medium (NHSM) (Gafni et al., 2013), but has been reformulated (www.stemcell.com). Our previous study revealed that RSeT-converted hPSCs display a transcriptomic profile reminiscent of primed hPSCs (Figure 1A) (Johnson et al., 2021). Subsequently, we derived RSeT hESCs by converting their primed counterparts using RSeT medium (Figure 1B) (Chen et al., 2024). In this study, we performed an unsupervised analysis of the top 200 significantly expressed genes based on our microarray data (Chen et al., 2024). Interestingly, we revealed a cluster of genes (e.g., *SREBF1*, *ACACA*, *FA2H*, *MTTP*, *AOPA1*, *ELOVL7*, and *GSTA5*) that are associated with metabolism (Figure 1C), suggesting that altered expression of metabolic genes might be a trigger of a global metabolite transformation that regulates pluripotent state transitions.

**Figure 1.**
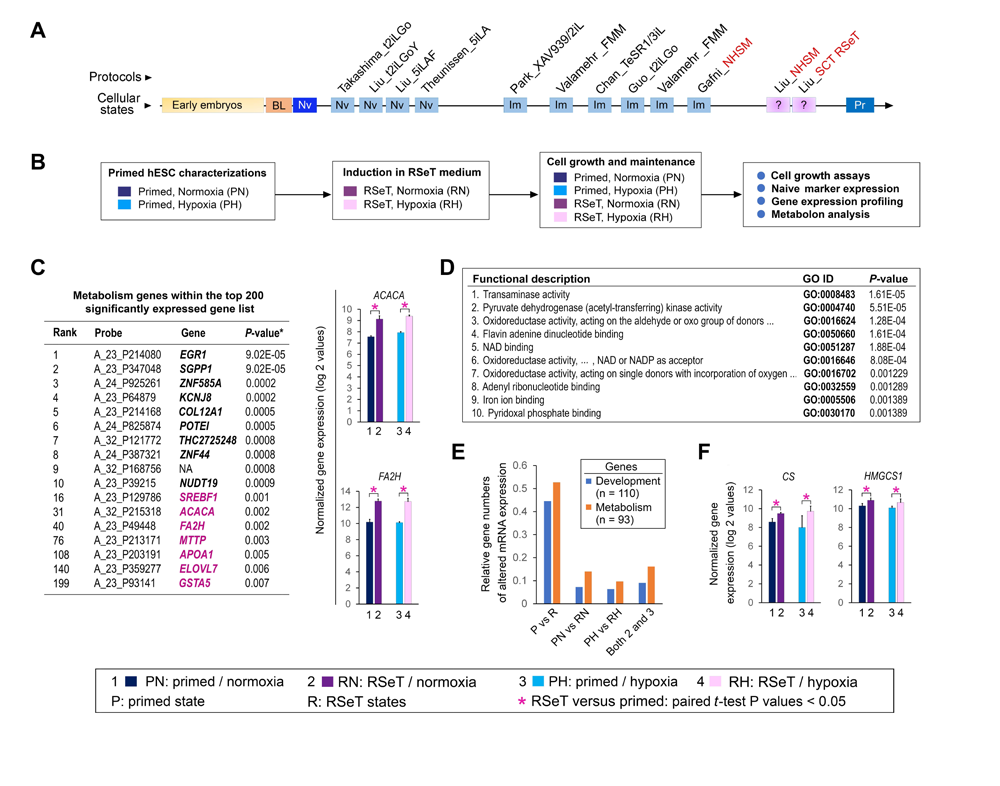
RSeT hESC line derivation and metabolic gene signatures. (**A**) Linear presentation of hPSCs from various naïve protocols as determined by meta-analysis (Johnson et al., 2021) under current naive conversion or reprogramming protocols (labeled in italicized characters with first authors’ names), t2iLGo, t2iLGoY, 5iLA, 5iLAF, XAV939/2iL, FMM, TeSR1/3iL, NHSM, and RSeT, from the laboratories of AS (Austin Smith), EZ (Elias Zambidis), HN (Huck-Hui Ng), JH (Jacob Hanna), JP (Jose Polo), PF (Peter Flynn), and RJ (Rudolf Jaenisch), highlighting the hPSCs (in red-font characters) that are related to RSeT protocols (SCT RSeT and NHSM). (**B**) Schema of hESC lines (H1, H7, and H9) treated under different primed and RSeT induction conditions as indicated. (**C**) Unsupervised gene signatures within the top 200 significantly expressed gene list, in which the genes associated with metabolism are highlighted in magenta color. *P*-values are type III ANOVA BH and FDR-corrected (Chen et al., 2024). Pair-wise analysis of the two representative metabolism genes (*ACACA* and *FA2H*), as depicted in the table of the left panel, expressed in primed and RSeT hESCs. (**D**) Ontological analysis of a supervised metabolism gene list (n = 93, Table S1), which is classified by 115 ontological categories (Table S2). Only the top 10 are shown with GO identification (ID) and adjusted *P* values. (**E**) Comparative analysis of relative gene numbers with altered mRNA expression in various pluripotent states. The complete gene lists are available in Table S1, which contains 93 genes associated with metabolism (n = 93), and in our recent report (Chen et al. 2024), which covers the genes known for development (n =112). (**F**) Pair-wise analysis of the two representative metabolism genes (*CS* and *HMGCS1*) in Figure 1E, expressed in primed and RSeT hESCs.

As shown in Figure 1D, ontological analysis of a supervised metabolism gene list (n = 93, Tables S1, S2) revealed top 10 functional categories, including transaminase activity to pyridoxal phosphate binding. Relative gene numbers with altered mRNA expression were generally higher in the metabolic gene group than those genes associated with development (Table S1, Figure 1E) (Chen et al., 2024). Thus, numerous metabolic genes (e.g., *SREBF1*, *ACACA, F2AH*, *ELOVL7*, *CS,* and *HMGCS1*) might be significantly upregulated in RSeT conditions (Figure 1F). With specific TaqMan qPCR probes, we confirmed that *SREBF1*, *ACACA*, *F2AH*, and *ELOVL7* were indeed significantly upregulated in RSeT hESCs or RSeT subsets, but not in primed hESCs cultured in hESC medium containing Knockout Serum Replacement (KSR) or mTeSR1 medium (Figure 2).

**Figure 2.**
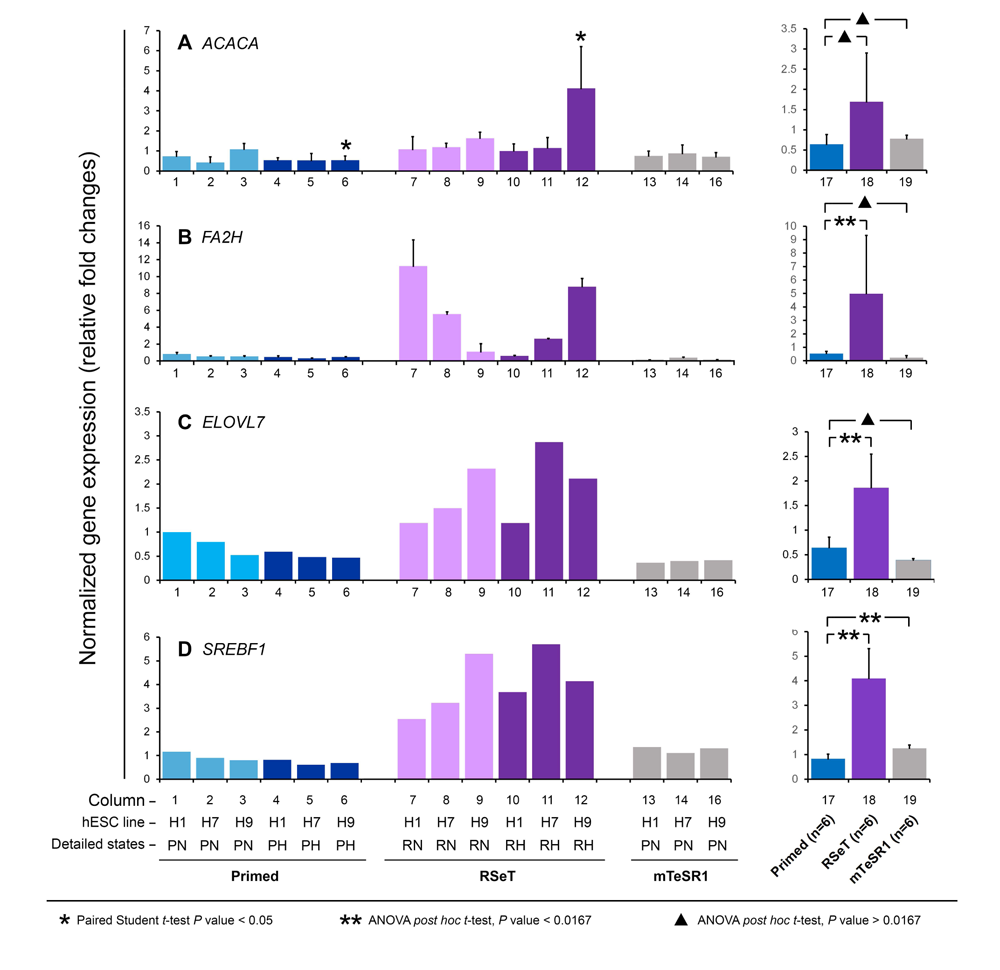
Quantitative real-time PCR (qPCR) confirmation of metabolic gene expression patterns identified from cDNA microarray (as shown in Figure 1C) in primed and RSeT hESCs as well as hESCs grown under mTeSR1. The right panel is the summary of data from individual cell lines under the three growth conditions shown in the left panel. Data are represented as the mean ± SD (standard deviation) with statistical analysis indicated.

These data revealed that metabolic genes are susceptible to RSeT-based growth conditions. We speculate that unscrambling extensively divergent metabolites in RSeT cells would provide robust reporter assays to assess the metabolic pathways and states, which ultimately determine specific cellular states and growth requirements for RSeT hPSCs.

### Globally mapping of metabolic changes in hypoxia versus normoxia and primed versus RSeT states

The present metabolic dataset comprises a total of 638 compounds of known identity (named biochemicals), which are involved in 9 super metabolic pathways (Figures 3A, 4A-C, Table S3). Following normalization to Bradford protein concentration, log transformation, imputation of missing values, Analysis of Variance (ANOVA) contrasts were used to identify 615 biochemicals that differed significantly between normoxia/hypoxia and primed/RSeT groups (Figure 3B). A summary of the numbers of biochemicals that achieve statistical significance (*P* ≤ 0.05), as well as those approaching significance (0.05 < *P* < 0.10), is shown in Figures 3B-D. Analysis by three-way ANOVA identified biochemicals exhibiting significant interaction and main effects for experimental parameters of hypoxia exposure, differential status, and cell line conversion (Figure 3B). For the comparison between primed and RSeT cells, the ratios of upregulated to downregulated metabolites increased 2.8-fold and 3.3-fold under normoxia and hypoxia (*P* < 0.05), respectively (see Figures 3C; 3E, left panel, 4B). However, for the comparison between hypoxia and normoxia, the ratios of up and down regulated metabolites in RSeT and primed hESCs are not significant (all *P* > 0.05) (Figures 3D; 3E, right panel; 4C). These data suggest that changes in oxygen levels have a much smaller impact on metabolic changes compared with the changes in cellular states.

**Figure 3.**
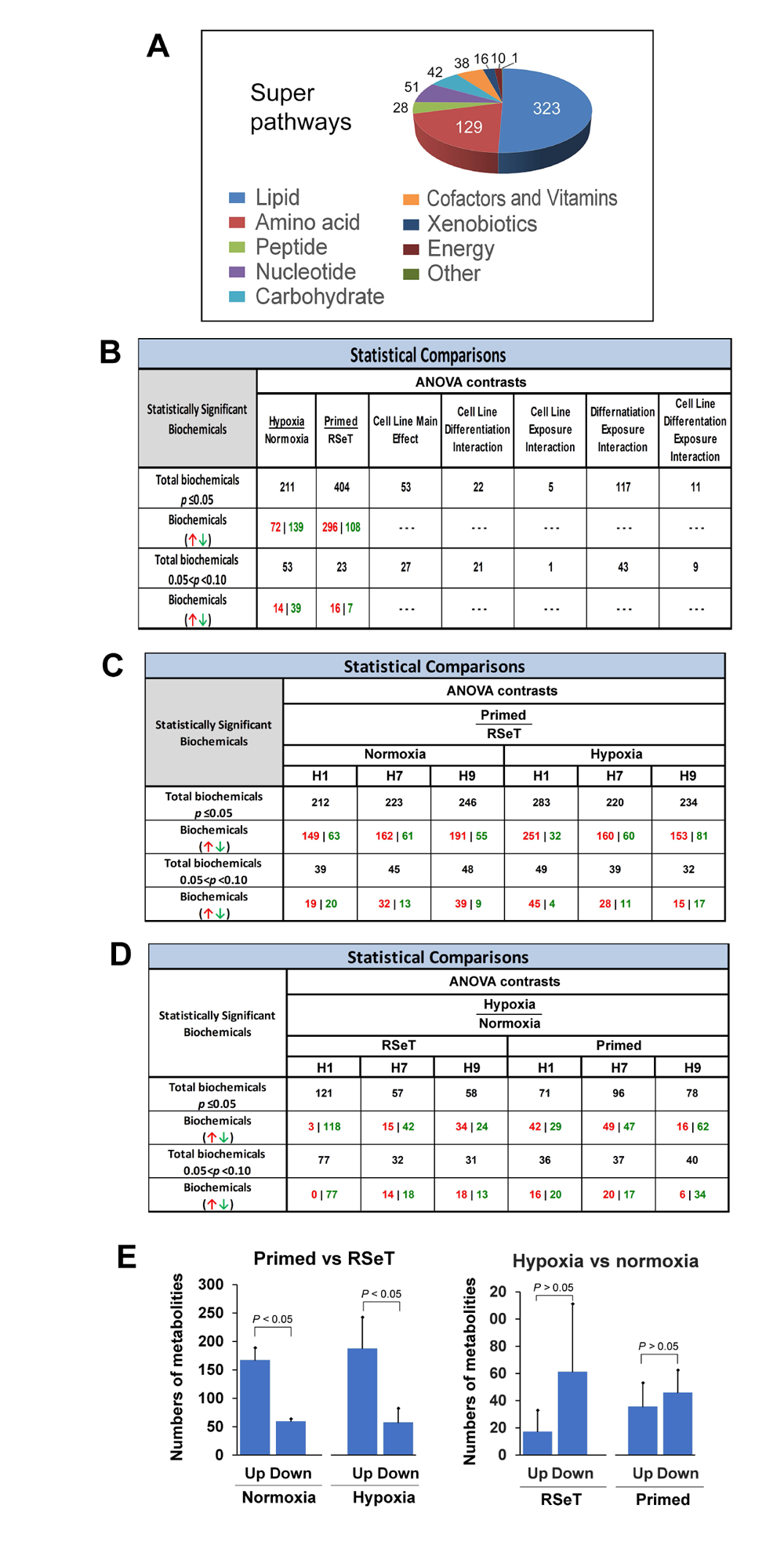
Global metabolic changes in primed versus RSeT hESC and hypoxia versus normoxia. (**A**) Metabolon super pathway summary. (**B**) Statistically significantly altered biochemicals. (**C** and **D**) ANOVA contrasts of primed versus RSeT and hypoxia versus normoxia comparisons under various conditions.

### RSeT hESCs retained active glycolysis but lack distinct glycolytic features as described in naïve hPSCs

Specifically, we employed the Metabolon platform (www.metabolon.com) to analyze glucose metabolism, including glycolysis, the tricarboxylic acid (TCA) cycle, and OXP (Figure 5A). Under hypoxic conditions, there was a significant increase in glucose uptake and glycolytic intermediates, suggesting an increased glucose availability and glycolysis, which were reflected in all comparisons by cell lines (though not significant in all *post-hoc* tests between primed and RSeT hPSCs) (Figures 5B, 5C1).

In general, RSeT hESCs did not show downregulated glycolysis (Figure 5B), as expected to occur as in the metabolic bivalency of naïve hPSCs. However, RSeT H9 cells exhibited the lowest glucose availability (Figure 5C1, lanes 9 and 12). Excess glucose can be converted to sorbitol or fructose. Both compounds (with sorbitol being represented by an isobaric contribution from both mannitol and sorbitol) tended to increase in RSeT cells especially in RSeT H9 cells, suggesting that glucose is in low glycolytic demand only in RSeT H9 cells (Figure 5C2).

A decrease in an isobar of sugar diphosphates (where fructose 2,6-bisphosphate is thought to be the dominant contributor) and the glycolytic intermediate, dihydroxyacetone phosphate (DHAP), in RSeT hESCs was consistent with decreased glycolytic use (Figure 5C4). Evidently, 3-carbon glycolytic intermediates 3-phosphoglycerate (Figure 5C5) and phosphoenolpyruvate (PEP) (Figure 5C6), downstream of DHAP, showed a similar metabolic pattern, only preferentially reduced under normoxic conditions (Figures 5C4-6). These data suggest a decrease in DHAP and an isobar of sugar diphosphates in RSeT cells could reflect an accelerating glycolytic flux toward non-pyruvate directions (Figures 5C3, 5C4).

However, pyruvate metabolism represents the last step of glycolysis and also the critical node to generate acetyl-CoA (for subsequent oxidation in the TCA cycle) and lactate that enables the regeneration of NAD^+^ to support high glycolytic rates (Figure 5A). Pyruvate in RSeT cells was significantly accumulated under normoxia, especially in RSeT H9 cells (Figure 5C7), suggesting decreased glycolytic/pyruvate metabolism. Overall, both pyruvate and lactate in RSeT hESCs remain at the same level as that of their primed counterparts. Moreover, pyruvate in RSeT cells was reduced to the same level of primed hESCs under hypoxia (Figure 5C7, lanes 10-12), which might suggest a rapid shift toward to the TCA cycle that supports oxidative phosphorylation (Figure 5A).

With respect to the TCA cycle, a decline in pyruvate could be represented by decreased acetyl-CoA (Figure 3C9), increased isocitrate (Figure 5C10), increased pool sizes of NAD^+^/NADH, and increased ATP production (Figures 3C15-18). We found no significant changes for three oxidative metabolites (i.e., succinate, fumarate, and malate) in RSeT hESCs (Figures 3C12-14), suggesting that the energetic outputs (reflected by ATP production) from the TCA cycle may be associated with the reductive glutamine metabolism that involves the conversion of α-ketoglutarate to citrate to fuel fatty acid (FA) synthesis (Figure 3A2) (Fendt et al., 2013; Metallo et al., 2011).

Taken together, the effects of oxygen tension on metabolic changes were limited when comparing primed with RSeT hESCs (Figures 4, S1). With respect to pluripotent states, changes in glycolysis itself are somewhat subtle. Nevertheless, our data suggest that RSeT hESCs retain a stable glycolytic state with enhanced OXP from non-TCA resources such as FAO. We anticipated that Seahorse XF metabolic assays would clarify the metabolic state in these RSeT cells.

**Figure 4.**
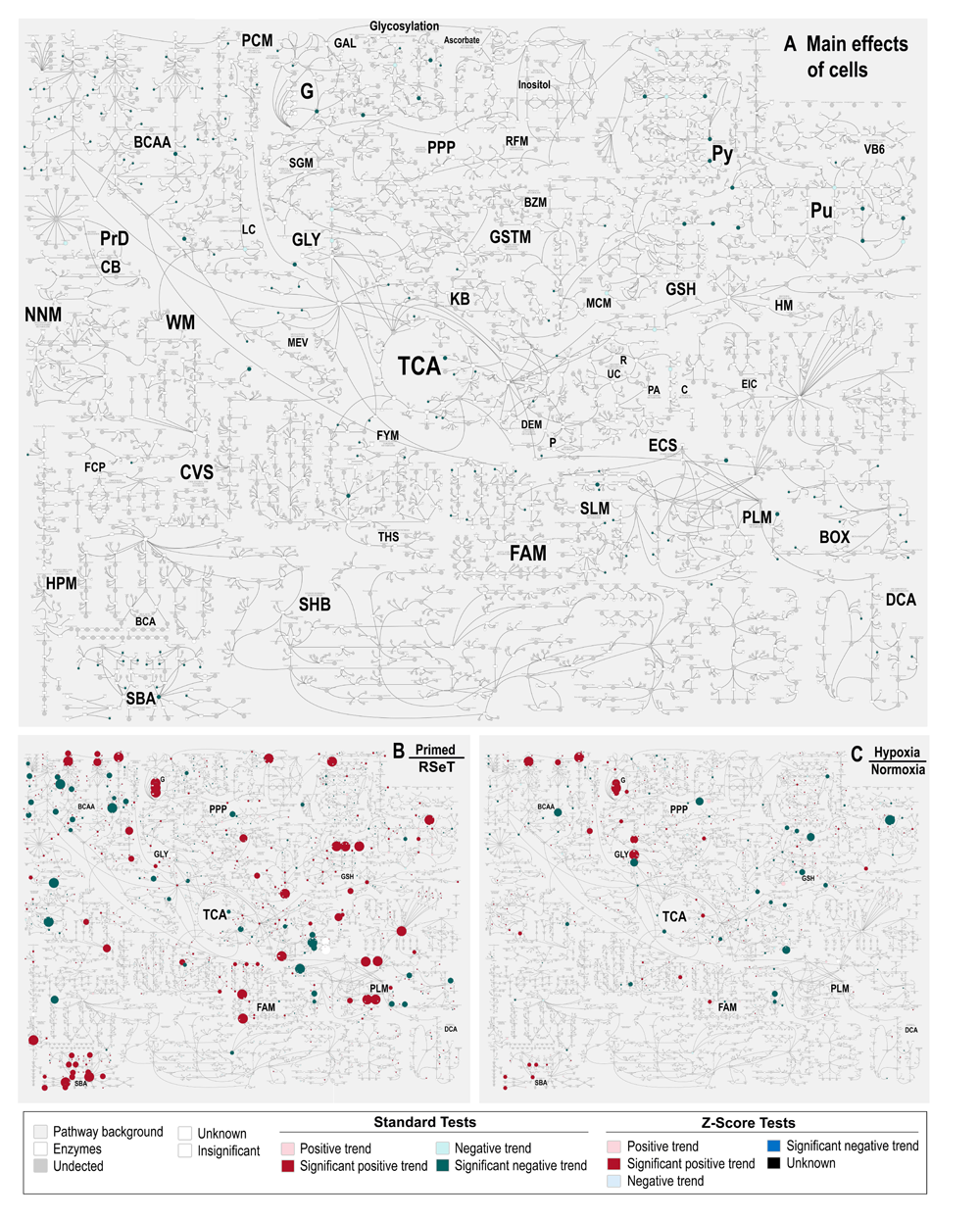
Globally mapping of metabolic pathway changes in primed versus RSeT hESC and hypoxia versus normoxia. Global view of the comprehensive Metabolon pathways (**A**), with individual metabolic pathways abbreviated and labeled in a larger font to facilitate a comparative analysis with (**B**) and (**C**), in primed and RSeT hESC lines (H1, H7, and H9, n = 3). A higher resolution of A is supplemented as Figure S1 to aid a rapid metabolic mapping.

### Seahorse glycolytic and mitochondrial oxidative phosphorylation assays support the metabolomic data

We performed the Seahorse XF Glycolysis Stress Test to verify the glycolytic changes described in Figure 5, via measuring the extracellular acidification rate (ECAR) in representative RSeT H9 cells (Figures 6A, 6B). Primed H9 cells grown on Matrigel in mTeSR1 were used as control for the analysis. Compared with primed cells, RSeT H9 cells showed significantly down-regulated glycolysis (*P* < 0.05), which was related to an elevated glycolytic reserve (Figure 6B). However, the glycolytic capacity remained at the similar level (*P* > 0.05) compared with primed control (Figure 6B). These data support the metabolomic finding that glycolysis remains globally unchanged in RSeT hESCs (Figures 5B, 5C).

**Figure 5.**
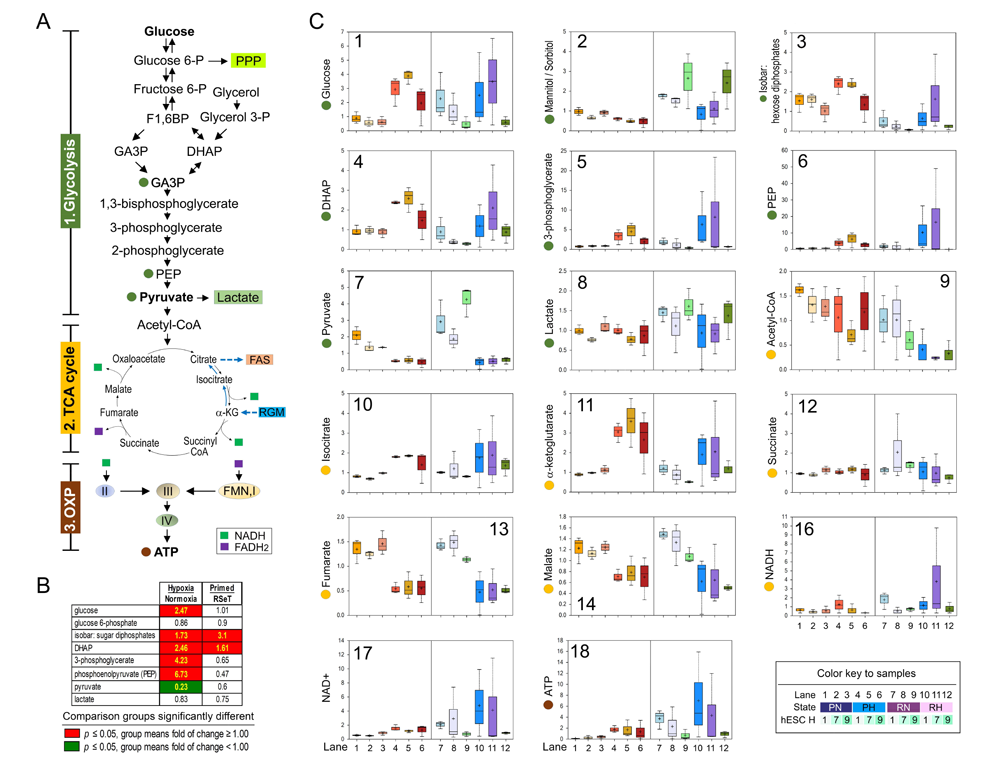
Glucose metabolism: glycolysis and oxidative phosphorylation in primed and RSeT hESCs. (**A**) Metabolic pathways of glucose to generate energy resources via glycolysis, tricarboxylic acid (TCA) cycle, and oxidative phosphorylation (OXP). (**B**) Representative metabolites of glycolysis, TCA cycle, and OXP as diagrammed in Figure 5A, with significantly altered mean metabolic values (*P* < 0.05) between samples (i.e., hypoxia versus normoxia and primed versus RSeT) highlighted with red or green colors in Figure 5B. (**C**) Quantitative analysis of metabolites related to glycolysis, TCA, and OXP as indicated in Figure 5A. Box plots denote the maximum, upper quartile, mean (+), median (the line that divides the box into two parts), lower quartile, and minimum values of three biological samples/replicates (n = 3) obtained from different passages from each cell line. There are 36 individual samples. Abbreviations: ATP, adenosine triphosphate; CoA, coenzyme A; DHAP, dihydroxyacetone phosphate; F1,6BP, fructose 1,6-bisphosphate; FADH2, dihydroflavine-adenine dinucleotide; GA3P, glyceraldehyde-3-phosphate; I, II, III, and IV, mitochondrial Complex I, II, III, and IV, respectively, of the electron transporter chain; NAD^+^, the oxidized form of nicotinamide adenine dinucleotide; NADH, 1,4-dihydronicotinamide adenine dinucleotide; P, phosphate; PEP, phosphoenolpyruvate; PPP, the pentose phosphate pathway; α-KG, alpha-ketoglutarate.

**Figure 6.**
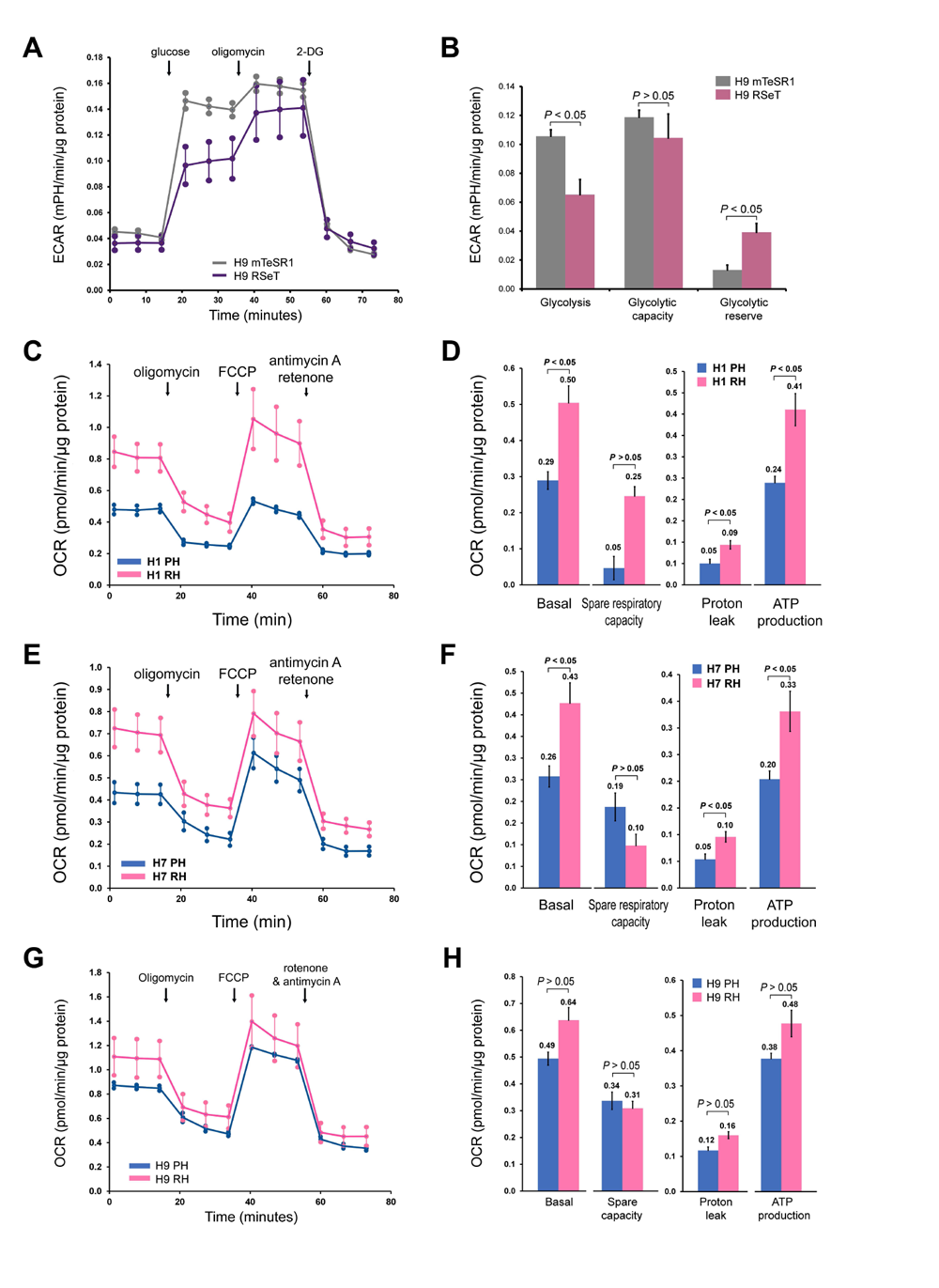
Agilent Seahorse XF metabolic functional assays in primed and RSeT hESCs. **(A** and **B).** The Seahorse XF Glycolysis Stress Test was used to assess the critical parameters of glycolytic function (i.e., extracellular acidification rate, ECAR). Consecutive compound injections facilitate the evaluation of glycolysis and glycolytic capacity, and glycolytic reserve in RSeT H9 cells. Primed H9 cells grown in mTeSR1 on Matrigel were used as control for the analysis. Both cell lines were cultured under normoxia. (**C-H**). The Mitochondrial Respiration XF Cell Mito Stress Test was carried out to measure the oxygen consumption rate (OCR) before and after the injection of oligomycin, FCCP, and a mix of rotenone and antimycin A as indicated **C**, **E**, **G**). Multiple parameters were calculated using Wave 2.6 Software (Agilent Technologies), including basal respiration, ATP Production, H^+^ (proton) leak, spare respiratory capacity, and nonmitochondrial respiration (**D**, **F**, **H**). OCR measurements were collected and normalized to protein content (as described above). *P* values were determined by two-tailed *t*-test. PH, primed hESCs maintained under hypoxia; RH, RSeT hESCs maintained under hypoxia.

We also carried out the Mitochondrial Respiration XF Cell Mito Stress Test to measure the oxygen consumption rate (OCR) as indicated (Figures 6C, 6E, 6G). Multiple parameters were calculated, including basal respiration, ATP Production, H^+^ (proton) leak, and spare respiratory capacity (Figures 6D, 6F, 6H). Among the three hESC lines, both RSeT H1 and H7 cells showed a significant increase in OCR, including 1.7-fold in basal OCR and 1.8-to 2-fold in protein leak, which was consistent with a 1.7-fold increase in ATP production in both lines (Figures 6C-F). However, RSeT H9 cells failed to show an elevated OCR (Figures 6G, 6H). These data are also consistent with the fact that RSeT H1 and H7 cell lines have higher ATP production than RSeT H9 cells as determined in metabolomics (Figure 5C18), possibly associated with oxidative phosphorylation contributed from additional energetic resources such as lipid metabolism.

### Robust lipid metabolic signatures that underlie the cellular remodeling capacities in primed and RSeT hESCs

To analyze the contribution of lipid metabolism to metabolic modes, we mapped all significantly altered metabolites into 70 pathways, delineating significant changes between primed and RSeT hESCs by principal component analysis (PCA) (Figure 7A). Interestingly, in the PCA plots, Principal component 1 (PC1) accounts for approximately 40% of the metabolic variability, only separating a handful of 4 hypoxic samples (e.g., H1 RH, H7 RH, H9 RH, and H9 PH) from the remaining groups. This separation could reflect the effects of experimental manipulations (e.g., the length of time exposed to hypoxic conditions in different passages), particularly in RSeT H1 cells under hypoxia (H1 RH). The major metabolic pathways that underlie the distinct pluripotent states are evident in PC2, which depicts 25% of the metabolic variations that clearly distinguish primed from RSeT hESCs, except H1 RH (Figure 7A). PC3 represents approximately 11% of metabolic variations, which separates normoxic from hypoxic conditions.

**Figure 7.**
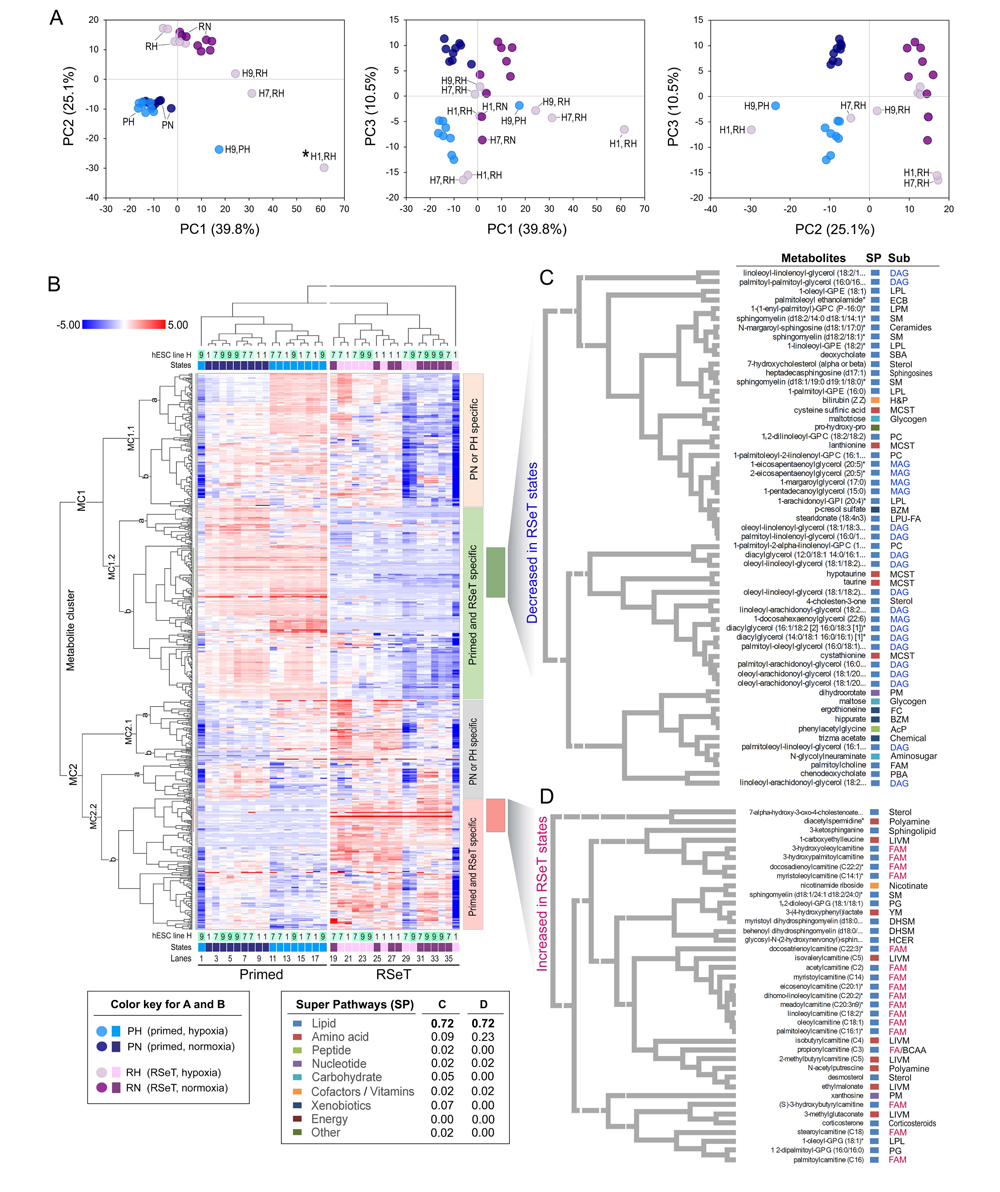
Principal component analysis (PCA) and metabolic hierarchical clustering analysis (HCA) of global metabolites associated with primed and RSeT hESCs. (**A**) PCA of metabolic signatures in hESCs with primed and RSeT states from 36 individual samples with three biological samples/replicates (n = 3) obtained from different passages from each cell line. Only major principal components (i.e., PC1, PC2, and PC3), which reveal 75% of metabolite variability, are shown. (**B**) Heatmap of the HCA cluster analysis of globally altered metabolites. (**C** and **D**) Enlarged representative view of decreased or increased metabolites in primed and RSeT hESCs, with metabolite descriptions, super pathways (SP, n = 9), and sub-pathways (Sub) labeled on the right panel. Abbreviations: AcP, acetylated peptides; BZM, benzoate metabolism; CER, ceramides; DAG, diacylglycerol; DHSM, dihydrosphingomyelins; ECB, endocannabinoid; FA/BCAA, fatty acid metabolism (also branched chain amino acid metabolism); FAM, fatty acid metabolism; FC, food component/plant; H&P, hemoglobin and porphyrin metabolism; HCER, hexosylceramides; LIVM, leucine, isoleucine and valine metabolism; LPL, lysophospholipid; LPM, lysoplasmalogen; LPU-FA, long chain polyunsaturated fatty acid (n3 and n6); MAG, monoacylglycerol; MCST, methionine, cysteine, SAM and taurine metabolism; PBA, primary bile acid metabolism; PC, phosphatidylcholine; PG, phosphatidylglycerol; PM, purine metabolism; SBA, primary bile acid metabolism; SM, sphingomyelins; YM, tyrosine metabolism.

Moreover, the major influential metabolites that emerged from the above PCA could be individually visualized in a high-resolution heatmap. We used a hierarchical clustering analysis (HCA)-based heatmap to assess the similarity between all samples (Figure 7B). In HCA, the first split in the dendrogram separated a single sample, H1 RH (Figure 7B, lane 36, also described as an outliner in Figure 7A). We present here 4 distinct subclusters that cover the altered metabolites that define primed and RSeT hESCs (Figure 7B). Clearly, these major clusters can differentiate normoxic from hypoxic conditions in primed hESCs (Figure 7B, lanes 1 to 18), but not in RSeT cells (Figure 7B, lanes 19 to 36), suggesting that heterogeneous responses occurred during the RSeT state conversion. Tertiary and quaternary splits in the dendrogram identified oxygenic exposure status, with surprisingly subtle effects associated with individual cell lines.

We also analyzed increased or decreased metabolites associated with RSeT cell lines in two representative subclusters. As indicated in Figure 7C, among the decreased metabolites (n = 57), 72% of them are lipids. Moreover, 45% (21/47) of these lipids were monoacylglycerol (MAG, n = 5) and diacylglycerol (DAG, n = 16) in RSeT lines. Noticeably, lysolipids [(e.g., 1-oleoyl-GPE 18:1 and 1-(enyl-palmitoyl)-GPC)], DAGs, and MAGs tended to decrease unidirectionally in RSeT cells in this study (Figure 7C, Table S3), suggesting that the stabilization of complex lipids led to decreased free FA availability in these RSeT cells.

Concurrently, among the increased metabolites (n = 39) in RSeT cells, 72% of them are also lipids, in which 61% of these lipids (principally acylcarnitines) are involved in FA metabolism, predominantly in FAO (Figure 7D). FAO occurs within the mitochondria, which is often de-emphasized in some stem cells due to an ‘immature’ morphology. The engagement of FAO in the mitochondria appears to be associated with both primed and RSeT hESCs (Figure 8).

**Figure 8.**
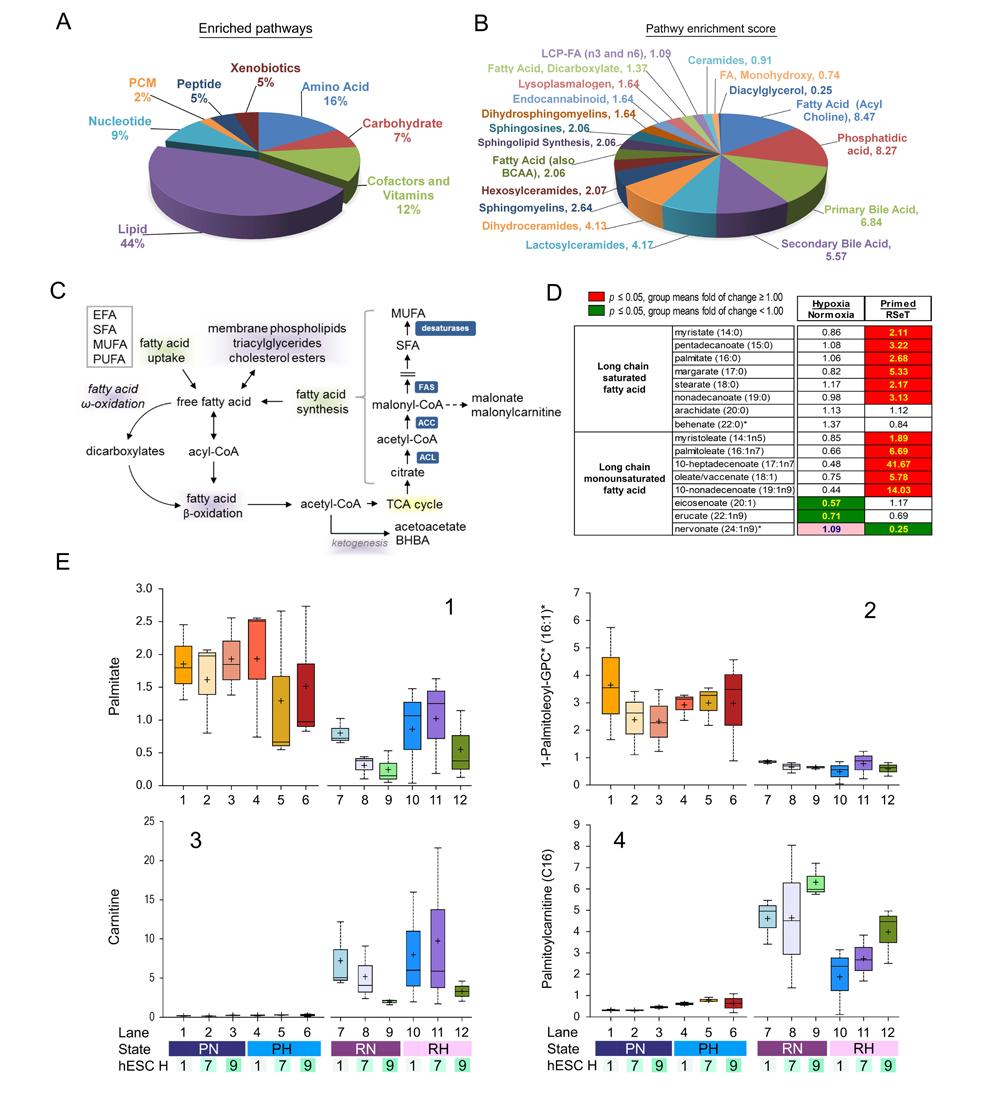
Metabolic pathway enrichment and lipid metabolism in primed and RSeT hESCs. (**A**) Pie graphic view of relative enrichment of 8 metabolic super pathways. (**B**) Lipid enrichment scores, ranging from 0.25 (diacylglycerol) to 8.47 (fatty acid metabolism, acyl choline) among 19 lipid sub-pathway categories (Table S3). (**C**) Schema of fatty acid biosynthesis and beta-oxidation pathways. (**D**) Changes of long chain saturated or monounsaturated fatty acid in hypoxia versus normoxia or primed versus RSeT hESCs, with significantly altered metabolites (P ≤ 0.05) highlighted in green and red colors. (**E**) Representative box plots of fatty acid metabolites in primed and RSeT hESCs, from 36 individual samples with three biological samples/replicates (n = 3) obtained from different passages from each cell line.

An unbiased metabolic pathway enrichment analysis also indicates that lipid metabolism ranks number 1 among 8 super pathways, constituting 44% of enriched pathways (Figure 8A). Breakdown of the lipid components shows FA acylcholine, phosphatidic acid, primary and secondary bile acids, lactosylceramides, dihydroceramides, sphingomyelins, hexosylceramides, and FA (also branched-chain amino acids) have the highest pathway enrichment scores, ranging from 8.5 to 2.1 (Figure 8B, Table S4). Relating to lipid metabolism, both FA biosynthesis and FAO are of particular interest because they are important resources of acetyl-CoA for energetic demands in addition to constituting membrane structures (Figure 8C).

### Increased FAO in RSeT hESCs

Consistently, hypoxia (compared to normoxia) showed few changes in lipid metabolites, suggesting that its impact on lipid oxidation was also not profound (Figure 8D, Table S3). In general, both long-chain saturated fatty acids (LC-SFA) (e.g., palmitate 16:0 and margarate 17:0) and long-chain monounsaturated fatty acids (LC-MUFAs) (e.g., 10-heptadecenoate 17:1n7 and 10-nonadecenoate 19:1n9) have relatively low presentation in RSeT cells (Figures 8D, 8E1, and 8E2), suggesting either a decrease in FA biosynthesis or an increase in FAO in these cells. Compared with primed hESCs, free FAs like palmitate (16:0) in RSeT hESCs were down-regulated (Figures 8D, 8E1), likely via an accelerated FAO of lipids derived from *de novo* FA biosynthesis as indicated by increased *ACACA* (known as *ACC* for biosynthesis) gene expression (Figure 8C; Figures 1C, 2A).

FAO is facilitated by a biochemical process called the carnitine shuttle, which involves the transesterification of a cytosolic carnitine molecule to the long-chain fatty acyl-coenzyme A (LCFA-CoA) to yield long chain acylcarnitines (LCACs). LCACs are imported into the mitochondria by carnitine-acylcarnitine translocase (CACT) and then re-esterified by CPT2 to generate LCFA-CoA that undergoes βIZoxidation (i.e., FAO) to produce acetyl-CoA. Acetyl-CoA is used to generate ATP via the TCA cycle and the mitochondrial electron transport chain (McCoin et al., 2015). Excess LCACs can be converted to ketone bodies (not detectable in our Metabolon data), acetyl-carnitine (increased in RSeT cells) or acetate (not detected in Metabolon data), retro-converted to LCACs and exported out of the mitochondria or the cells (Table S3). Drastically increased free carnitine in RSeT cells was apparently associated with elevated LCACs and the higher demand in FAO (Figures 8E2, 8E3). The levels of accumulated LCACs between individual RSeT cell lines reflect the differences in FAO rates and the flux of acetyl-CoA into the TCA cycle. Both RSeT H1 and H7 cell lines seem to have less accumulated LCACs (Figure 8E4), conceivably associated with higher ATP production than RSeT H9 cells (Figures 5C18, 6C-H). Nonetheless, besides LCACs, a large body of enriched metabolites in this study might be used to identify cell-line-specific metabolic signatures, indicative of cellular states.

### Globally enriched cell-line-specific metabolic signatures in primed and RSeT hESCs

Due to the differential proliferative capacity of hESCs under RSeT conditions (Chen et al., 2024), it is imperative to address the metabolic linkage associated with unfavorable cell growth in some RSeT cells (e.g., RSeT H1 cells). Accordingly, we defined metabolic signatures by classifying significantly altered metabolites (n = 432) into 36 groups (G1 to G36) (Figure 9).

**Figure 9.**
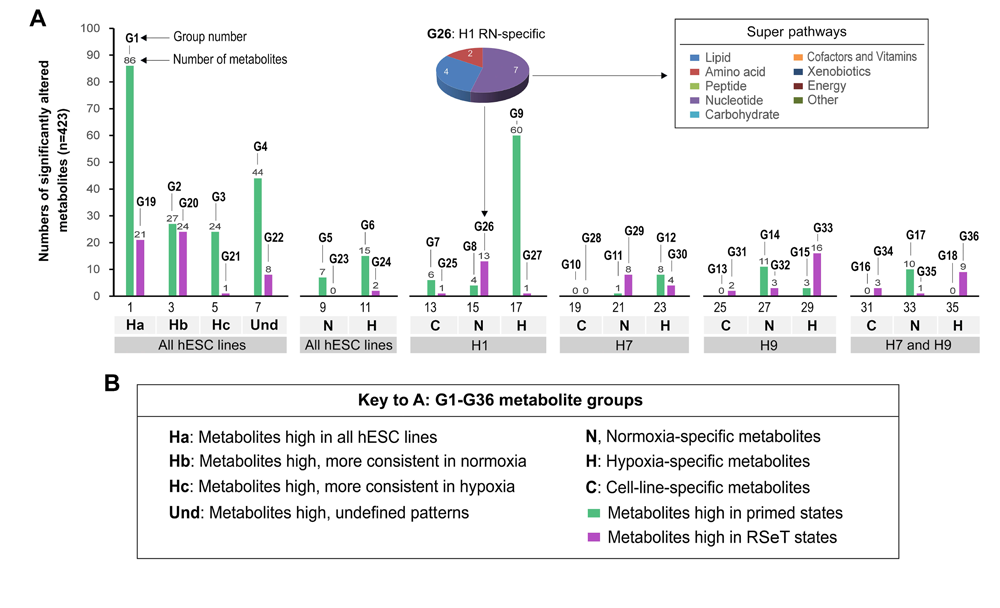
Globally enriched cell-line-specific metabolic signatures in primed and RSeT hESCs. (**A**) Metabolic signatures underlying cell-type-and pluripotent-state-specific conditions in primed and RSeT hESCs. Numbers of significantly altered metabolites (n = 423) are classified into 36 groups (G1-36, Table S3), which are used to define hESC-line-specific and pluripotent state-specific conditions as detailed in the bottom panel (**B**). Inset of Figure 9A: representative pie graphic view of the numbers of metabolic alterations in primed and RSeT hESCs.

Briefly, group 1 (G1) delineates metabolites (n = 86) high in all primed hESCs, comprising lipid (72%), amino acid (13%), nucleotide (1%), carbohydrate (3%), cofactors and vitamins (1%), peptide (5%), and xenobiotics (5%). In contrast, G19 has only 21 metabolites highly observed high in all RSeT cell lines, containing lipid (71%), amino acid (14%), cofactors and vitamins (14%) (Figure 9A, column 2). Thus, compared with primed hESCs, a sharp decrease in metabolites with a relatively 14-fold elevated percentage of cofactors and vitamins represents one of the major metabolic features in RSeT cells. Interestingly, G5 depicts PN (primed-normoxic)-specific metabolites (n = 7), which are composed of 100% lipids (including adrenate, MAG, DAG, and propionyl CoA), whereas G6 covers PH (primed-hypoxic)-specific metabolites (n = 15), displaying 40% of lipids concomitantly with a 47% of amino acid metabolites (Figure 9A, column 11). These data suggest that the balance between lipid and amino acid metabolism may be important for the maintenance of primed states and hPSC homeostasis under hypoxia.

Also interestingly, G7-9, G12, G14, and G15 revealed cell-line-specific metabolites of primed hESC lines, in which 6 metabolites (e.g., inosine, tricosanoyl sphingomyelin, and 5’-GMP) in G7 characterize H1 cells, independent of oxygen tension. Compared with G8 (H1 PN-specific metabolites, n = 4), G9 had a 15-fold increase in H1 PH-specific metabolite numbers (n = 60), the highest number of metabolites among the 3 hESC lines (Figure 9A, column 17). G9 constitutes lipid (52%), amino acid (30%), nucleotide (5%), cofactors and vitamins (5%), carbohydrate (3%), and peptide (3%) metabolites. However, the potential functional implications in H1 cells remain to be determined.

Likewise, the H7-and H9-specific metabolites may be related to their survival and proliferative capacities under the RSeT conditions. Regarding H7 cells, there is a significant increase in leucylglutamine, methylphosphate, 7-alpha-hydroxy-3-oxo-4-cholestenoate, ophthalmate, leucylglycine, sphingomyelin, palmitoyl dihydrosphingomyelin, methyl glucopyranoside (alpha and beta) in RSeT H7 cells under normoxia (H7 RN) (Figure 9A, G29; Table S3). Additionally, 4 metabolites that include diacetylspermidine, arabitol (xylitol), docosatrienoate (22:3n6), and kynurenate were found high in H7 RH (Figure 9A, G30; Table S3).

Concerning RSeT H9 cells, there is a 3.3-fold increase in 7-methylguanine, pyruvate, and urate under H9 RN (Figure 9A, G32; Table S3). Correspondingly, a panel of metabolites were significantly elevated under H9 RH, which again include diverse lipids (1-palmitoyl-2-oleoyl-GPG, lactosyl-N-nervonoyl-sphingosine, 1-stearoyl-2-oleoyl-GPG, 1,2-dioleoyl-GPI, stearoyl-carnitine, 1-palmitoyl-2-oleoyl-GPA, and N-erucoyl-sphingosine), the neurotransmitter 4-hydroxybutyrate, mannitol/sorbitol, N-formyl-methionine, pyridoxal phosphate, 2-oxoarginine, 5-methyluridine, 4-hydroxyphenylpyruvate. 2-methylcitrate (homocitrate), and citrulline (Figure 9A, G33; Table S3). However, the exact roles of these increased metabolites in RSeT H7 and H9 cells remain to be determined.

Nevertheless, defining metabolic signatures in H1 cells is also of particular interest because of their retarded cell growth rates under the RSeT growth conditions (Chen et al., 2024). Noteworthy, there is a 2.3-, 2.5-, 2.4-, 3.2-, and 1.7-fold increase in UMP, AMP, N-acetylglutamine, phenylpyruvate, and N-acetylserine in primed H1 cells, respectively, compared with H1 RH (Figure 10: G7, G9; Table S3). These data highlight that nucleotide metabolites (e.g., UMP and AMP), which are under-investigated in hPSCs, might be essential for hESC (e.g., H1 cell) growth and metabolic plasticity.

**Figure 10.**
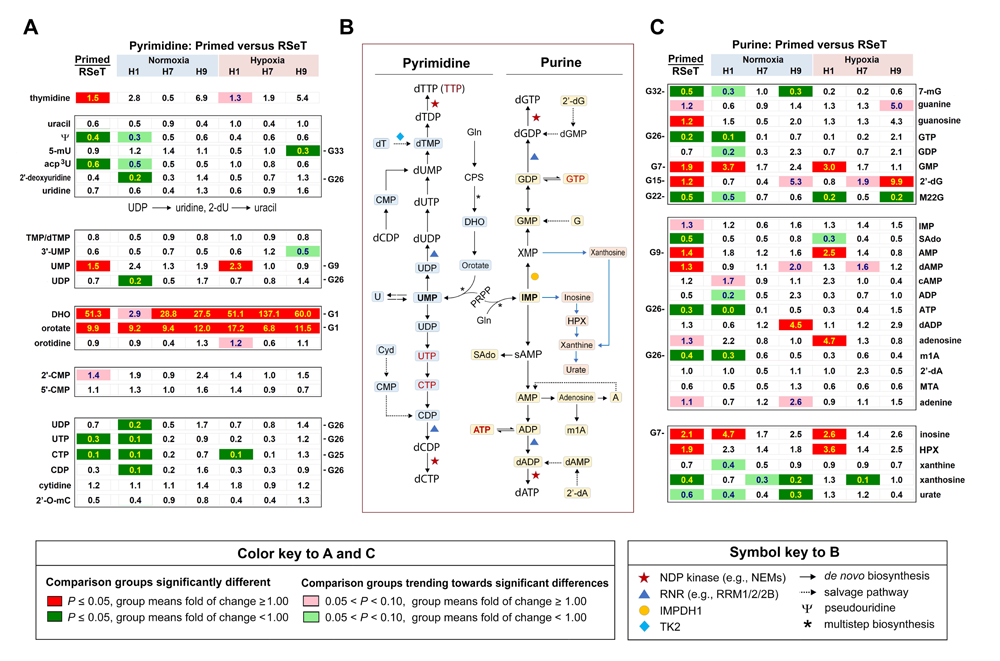
Nucleotide metabolism associated with purine and pyrimidine pathways. Representative pairwise comparison of metabolites of pyrimidine (**A**) and purine (**C**) metabolism of primed versus RSeT hESCs (under hypoxia versus normoxia) in 36 individual samples with three biological samples/replicates (n = 3) from each cell line. Significantly altered metabolites (*P* ≤ 0.05) are highlighted in green and red colors. (**B**) The diagram of the purine and pyrimidine metabolic pathways with symbol keys in the bottom panel.

### Imbalanced nucleotide and nucleoside metabolism

Fortunately, we were also able to identify several notable signatures enriched with nucleotide or nucleoside metabolism (Figures 10A-C, S2). First, we found relatively low levels of dihydroorotate and orotate in RSeT cells, indicating significantly decreased *de novo* pyrimidine biosynthesis in these cells (Figure 10A). Second, lower levels of AMP, GMP, and inosine, particularly in H1 cells, may also suggest decreased *de novo* purine biosynthesis in RSeT cells (Figure 10C). Likely, RSeT cells may utilize pyrimidine salvage pathways (via thymidine, uridine, uracil, and cytidine) for DNA or RNA synthesis (Figure 10B).

Furthermore, there were a total of 11 metabolites significantly increased in RSeT H1 cells in a H1-cell-line-, H1-RN-, and H1-RH-specific manner. For example, G26 describes H1-RN-specific metabolites (n = 13), which contain reduced lipid (31%) and amino acid (15%) presentation, concomitantly with an expressively increased nucleotide metabolite by 54% (n = 7) (Figures 9A, 10). Particularly, there was a 10-fold increase in CTP in both H1 RN and RH, a more than 10-fold increase in GTP, ATP, UTP, and CDP in H1 RN (Figure 10), and a 5-fold increase in both UDP and 2’ deoxyuridine in H1 RN (Figure 10A: G25, G26).

Collectively, these data imply that the accumulation of NTPs (e.g., ATP), not dNTPs, in the nucleotide pool in RSeT H1 cells, may inhibit DNA synthesis. Also, a significant decrease in AMP:ATP and UMP:UTP ratios in RSeT H1 cells (Figure 10, Table S3) might account for the energetic status that responds to the metabolic stress and autophagy. The significantly elevated level of S-(1,2-dicarboxyethyl) glutathione (DCEG) (Figure 9, G27; Table S3) was an indicator of the response to higher reactive oxygen species (ROS) stress associated with nucleotide metabolism in RSeT H1 cells.

In summary, the results from global metabolomic profiling and metabolic functional assays highlight several important changes related to exposure to hypoxia and to pluripotent state transitions. In general, hESCs exposed to hypoxic conditions showed distinct features that are associated with the increased use of glycolysis. However, the hypoxic effects on global gene expression and metabolites, especially lipids, are limited. RSeT hESCs exhibit primed hESC-like transcriptomes with notable metabolomic features, consistent with the activation of mitochondrial metabolism by increased FAO as well as elevated metabolic stress in a cell-line-specific manner.

## DISCUSSION

Adequate regulation of metabolic requirements is crucial in achieving optimal reprogramming efficiency, as it controls the growth, self-renewal, differentiation of hPSCs, and promotes the ability of somatic cells to undergo reprogramming. Yet, different types of cells utilize diverse metabolic modes to implement their cellular activities. It has also been established that primed hPSCs exclusively use glycolysis (i.e., monovalency), whereas naïve hPSCs utilize a bivalent mode. Regarding RSeT hPSCs, it is clear that these cells develop a metabolic multiplicity (i.e., quadrivalency), demonstrated by several noteworthy characteristics that will be elaborated below.

A first notable metabolic feature in RSeT hESCs is that the hypoxic conditions have much less effect on global metabolism in these hESC with a partially primed pluripotent (PPP) state (Chen et al., 2024), compared with pluripotent state conversion in these cells (Figures 3, 4). Also, the activation (or stabilization) of HIFs by hypoxic conditions only leads to alterations in glucose utilization with subtle changes in glycolysis (Figure 5). This characteristic could potentially allow the RSeT cells to thrive and passage in both hypoxic and normoxic conditions. Consistently, a similar formulation of RSeT medium, termed NHSM, can also support hPSC growth and maintenance independent of oxygen tension (Gafni et al., 2013). Previous reports suggest that hypoxic conditions change both metabolism and epigenetic marks in hPSCs (Lees et al., 2019; Spyrou et al., 2019). Human naïve protocols generally require hypoxic conditions, which allow HIFs to regulate pluripotent states and cellular proliferation (Forristal et al., 2010; Somasundaram et al., 2020; Tsogtbaatar et al., 2020). Thus, this metabolic feature provides, at least, a partial resolution of the inconsistency of oxygen tension requirements in hPSC culture.

The second remarkable metabolic feature of RSeT hESCs is the activation of mitochondrial metabolism, which is associated with active lipid metabolism (including elevated LCACs and high FAO demand) rather than the TCA cycle (Figures 5-8, Tables S3, S4). It is known that LCACs modulate insulin sensitivity, PKC signaling, cellular stress response, and plasma membrane interactions in some cellular models. High concentrations of LCACs may lead to the activation of cellular stress pathways (MAPK, JNK, ERK and p38 MAPK) and caspaseIZ3 (McCoin et al., 2015). Thus, LCACs in RSeT cells might hinder the primed-to-naïve transition through enhancing MAPK, ERK, and PKC activity required for stabilizing the primed pluripotent state. As a result, RSeT hESCs acquire a unique state downstream of formative pluripotency (Figure 11). Similarly, primed hPSCs could be converted to a stable primed-to-naïve intermediate in lipid-free conditions, likely *via* activated *de novo* lipid biosynthesis and the inhibition of endogenous ERK signals (Cornacchia et al., 2019).

**Figure 11.**
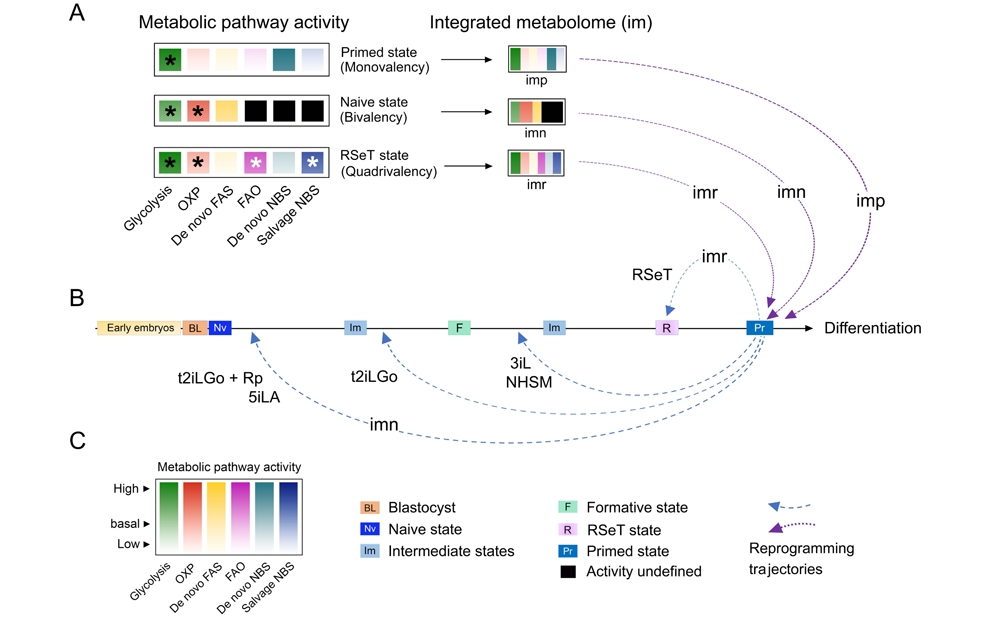
Regulation of pluripotent states by integrated metabolome inputs. (**A**) Regulation of primed, naïve, and RSeT-based pluripotent states by monovalent, bivalent, and quadrivalent metabolic modes, respectively, as indicated by color-coded metabolic activity and asterisk signs. Multimodal metabolic pathways function as individually integrated metabolomes (im), in which imp, imn, and imr represent individual metabolome for primed, naïve, and RSeT-based states, respectively. (**B**) Integrated metabolomes for the maintenance of primed hPSCs and/or reprogramming the primed state to naïve, formative, and RSeT-based states. Differential metabolic capacity may limit the reprogramming efficiency and trajectories observed in different naïve conversion protocols [such as t2iLGo plus reprogramming transgenes (t2iLGo + Rp), 5iLA, t2iLGo, 3iL, NHSM, and RSeT], resulting in diverse pluripotent states upstream or downstream of the formative state. (**C**) Color key to metabolic pathway activity (depicted in color gradients) and abbreviations or symbols to cell types, cellular or pluripotent states, and reprogramming trajectories. Additional abbreviations are: FAS, fatty acid biosynthesis; NBS, nucleotide or nucleoside biosynthesis; OXP, oxidative phosphorylation.

The third significant feature of RSeT hESCs is the dysregulated nucleotide metabolism, leading to cellular stress, lower metabolic reprogramming efficiency, and retarded cell growth or cell death in these cells (Figures 10, S2). Striking nucleotide metabolic landscapes include significantly increased CTP, GTP, UTP, and ATP, which may be associated with the adverse cell growth in RSeT H1 cells (Figures 10, 11). Moreover, the imbalanced presentation of dNTPs without the presence of dTTP may also have detrimental effects on DNA synthesis, thus deterring cell replication or growth (Diehl et al., 2022; Mathews, 2015). In addition, the decreased AMP:ATP ratio might reduce the capacity of RSeT H1 cells to activate the AMPK pathway that confers the cells resistant to autophagic cell death signals (Chen and Calderone, 2016; Rubinsztein et al., 2011). Hence, the low AMP:ATP ratio may be required for RSeT H1 cells to survive at the cost of generating excessive reactive oxygen species (ROS). Thus, metabolic changes in both primed and RSeT H1 cells provide a rational basis for optimizing hPSC cellular state and growth with physiologically relevant metabolic modalities (Figure 11A).

Regarding metabolic modalities, it is still not well understood what the optimal metabolic mode is and how individual hPSC lines utilize it in a specific pluripotent state. As introduced in the introduction section, naïve pluripotent stem cells prefer a bivalent metabolic mode (i.e., glycolysis and OXP) whereas primed hPSCs use an almost exclusive glycolytic mode (Takashima et al., 2014; Theunissen et al., 2014; Zhou et al., 2012). Our data reveal that RSeT cells employ a unique lipid metabolic mechanism, which couples fatty acid biosynthesis with FAO in addition to glycolysis, OXP, and imbalanced nucleotide metabolism, thereby constituting a distinct quadrivalent mode. This quadrivalency has not been reported previously in naive hPSCs, which likely limits RSeT cell’s reprogramming capacity toward formative and naive pluripotency (Figure 11).

In conclusion. we have redefined a unique pluripotent state between formative and primed pluripotency in RSeT hESCs in terms of their metabolic traits. RSeT cells are relatively insensitive to hypoxic tension, exhibiting a quadrivalent metabolic mode (including an enhanced FAO and imbalanced nucleotide metabolism). Primed hPSCs may have a restricted metabolic plasticity to be converted to the formative and naïve states under certain hPSC growth conditions (such as RSeT). Our study suggests that a physiologically compatible metabolic mode may be instrumental for hPSC growth and their broad downstream applications.

## MATERIALS AND METHODS

### Primed hESC culture on mouse embryonic fibroblast (MEF) feeder cell layers

All hESC lines (H1, H7, and H9) were cultured on MEF feeder cell layers in hESC medium, based on DMEM/F12 medium and Knock-Out Serum Replacement (KSR), in 6-well polystyrene plates. These cells were maintained in a 5% CO_2_ incubator at 37°C and supplemented with 3% (hypoxia) or 20% O2 (normoxia). Detailed information about hESC passage, expansion, and cryopreservation is available in our previous reports (Chen et al., 2024; Chen et al., 2012; Mallon et al., 2013).

### RSeT hESC culture

The three hESC lines H1, H7, and H9 were converted to a different pluripotent sate using the commercially available RSeT™ medium (www.stemcell.com). RSeT hESCs were cultivated and maintained under the same conditions as their primed counterparts as described above and detailed in our recent report (Chen et al., 2024).

### Quantitative real-time PCR (qRT-PCR)

For qRT-PCR, genomic DNA-free RNAs (2 μg) were reversely transcribed (in a 20-μl volume) into cDNAs. All TaqMan™ MGB probes were purchased from Thermo Fisher Scientific Inc. (see Key Resource Table). Both β-actin (*ATCB*) and glyceraldehyde-3-phosphate dehydrogenase (*GAPDH*) assays were used as internal controls for normalization. Approximately, 100 ng of cDNAs were amplified in 1X TaqMan™ Fast Advanced Master Mix and 1X TaqMan™ Assay in a 20-mL reaction using the QuantStudio 6 Flex real-time PCR Platform (Thermo Fisher Scientific Inc.) according to the manufacturer’s instructions. Data were analyzed by a three-step comparative cycle threshold (Ct) method described previously (Chen et al., 2024).

### The Metabolon platform for global metabolites profiling

We analyzed primed (n = 18) and RSeT (n = 18) hESC samples employing the precision metabolomics from Metabolon Inc. (Morrisville, NC), which has more than 3,300 commercially purified standard compounds in the proprietary reference library that elucidates 70 metabolic pathways. We unbiasedly analyzed all 638 metabolites in 8 super pathways, which include lipid (50%), amino acid (20%), nucleotide (8%), carbohydrate (7%), cofactors and vitamins (6%), peptide (4%), xenobiotics (3%), and energy (2%). The following processes were implemented to maximize the data reliability and reproducibility, which include: (i) controlling the instrument and process variability, sample accessioning and preparations, and quality assurance/quality control (QA/QC). Ultra-high-performance liquid chromatography-tandem mass spectroscopy (UPLC-MS/MS) was used for the analysis. The detailed methods are described as follows.

#### Instrument and process variability

Preceding the injection of samples into the mass spectrometers, the internal standards were included in all samples to determine the median relative standard deviation (RSD) for the instrument variability. The median RSD of all endogenous metabolites (as non-instrumental standards) of all experimental “Matrix” samples, which are technical replicates of pooled experimental samples, was calculated to assess the overall process variability. The values of both instrumental and processing variabilities met Metabolon’s acceptance criteria.

#### Sample preparations

Primed and RSeT hESC colonies were dissociated from MEF layers, collected in 1.5 mL Eppendorf tubes, snap-frozen into liquid nitrogen, and immediately stored in a −80°C freezer prior to sample processing. All samples were accessioned into the Laboratory Information Management System (LIMS) and assigned by the Laboratory Information Management System (LIMS) with a unique identifier. Samples were prepared using the automated MicroLab STAR^®^ system from Hamilton Company. Prior to the first step in the extraction process, recovery standards were added for QC purposes. To recover chemically diverse metabolites that are associated with protein or protein matrices, samples were precipitated by methanol with 2-minute vigorous shaking (Glen Mills GenoGrinder 2000) and then centrifugated. The organic solvent in the sample extracts was removed using TurboVap® (Zymark). The sample extracts were stored overnight in liquid nitrogen prior to the analysis. The final extract of each sample, divided into five portions, was used for the analysis by two separate reverse-phase (RP)/UPLC-MS/MS methods with positive (n = 2) or negative (n =1) ion mode electrospray ionization (ESI), and HILIC/UPLC-MS/MS with negative ion mode ESI (n = 1). One sample was used as a backup.

#### QA/QC

Jointly with the experimental samples, different types of controls were also used in the analysis. These controls include (i) extracted water samples as process blank, (ii) a cocktail of QC standards, and (iii) a technical replicate generated from a pooled experimental matrix by taking a small aliquot of each experimental sample. Experimental samples were randomized across the platform run with QC samples spaced evenly among the injections. The overall process and platform variability can be estimated based on the variability among consistently detected biochemicals.

### UPLC-MS/MS

All methods employed a Waters ACQUITY ultra-performance liquid chromatography (UPLC) and a Thermo Scientific Q-Exactive high resolution/accurate mass spectrometer. These mass spectrometers are interfaced with a heated electrospray ionization (HESI-II) source and a 35,000-mass-resolution Orbitrap mass analyzer. To safeguard the injection and chromatographic uniformity, the sample extract was dried and reconstituted in solvents containing a series of standards at constant concentrations. One aliquot was examined using acidic positive ion conditions, chromatographically optimized for more hydrophilic compounds. In this case, the extract was gradient eluted from a C18 column (Waters UPLC BEH C18-2.1 x 100 mm, 1.7 µm) using water and methanol, containing 0.05% perfluoropentanoic acid and 0.1% formic acid. The second aliquot was also evaluated using acidic positive ion conditions but optimized for more hydrophobic compounds. In this process, the extract was gradient eluted from the same C18 column described above, with methanol, acetonitrile, water, 0.05% perfluoropentanoic acid, and 0.01% formic acid. The third aliquot was analyzed using a basic negative ion optimized condition with a dedicated C18 gradient column, in which basic extracts were eluted using methanol and water containing 6.5 mM ammonium bicarbonate (pH 8.0). The fourth aliquot was eluted from an HILIC column (Waters UPLC BEH Amide 2.1 x 150 mm, 1.7 µm) via negative ionization and a gradient consisting of water and acetonitrile in the presence of 10 mM ammonium formate (pH 10.8). Raw data files are archived for further analysis.

### Seahorse XF Glycolysis Stress Test

The assay was performed with an Agilent Kit 103020-100 (Agilent Technologies). One day prior to assay, a sensor cartridge was hydrated in XF calibrant at 37°C in a non-CO_2_ incubator overnight. Primed and RSeT hESCs were plated as small clumps at a pre-determined density of 1 to 3 x 10^5^ cells per well in a 96-well Seahorse XF Microplate coated with 5% Matrigel. At the day of assay, stem cell medium was replaced with Seahorse XF DMEM assay medium (pH 7.4) containing 2 mM glutamine. The cells were incubated at 37°C in a non-CO_2_ incubator for 60 minutes. Before starting the XF assay, the assay medium was replaced with warm assay medium again. Meanwhile, freshly made 10X compounds were loaded into the injection ports as follows: Port A (100 mM glucose), Port B (10 μM oligomycin), and Port C (500 mM 2-DG) in the XFe/XF96 sensor cartridge. The cell plate was loaded into the Seahorse XFe/XF96 Analyzer. Extracellular acidification rate (ECAR) measurements were collected and normalized to protein content that was determined by Pierce™ BCA Protein Assay (Sigma, Catalog number 23228). Glycolysis, glycolytic capacity, and glycolytic reserve were calculated by Wave 2.6 Software (Agilent Technologies).

### Seahorse XF Mito Stress Test

The assay was performed with Agilent Kit 103015-100 (Agilent Technologies), following the same procedures of preparing the sensor cartridge and of cell seeding. At the day of assay, stem cell medium was replaced with Seahorse XF DMEM assay medium (pH 7.4) containing 20 mM glucose, 2 mM pyruvate, and 2 mM glutamine. The cells were incubated at 37°C in a non-CO_2_ incubator for 60 minutes. Before starting the XF assay, the assay medium was replaced with warm assay medium. Meanwhile, freshly made 10X compounds were loaded into the injection ports as follows: Port A (15 μM oligomycin), Port B [5 μM carbonyl cyanide 4-(trifluoromethoxy)-phenylhydrazone (FCCP)], and Port C (10 μM retenone/antimycin A) in the XFe/XF96 sensor cartridge. The cell plate was loaded into the Seahorse XFe/XF96 Analyzer. Oxygen consumption rate (OCR) measurements were collected and normalized to protein content (as described above). The basal oxidative phosphorylation rate, spare respiratory capacity, proton leak, and ATP production were calculated by Wave 2.6 Software (Agilent Technologies).

### Bioinformatics of Metabolon datasets

The LAN backbone and a data server (running Oracle 10.2.0.1 Enterprise Edition) serve the foundations for the software and hardware that support bioinformatic analysis. The analysis utilizes multiple modules that include the LIMS and tools for data extraction, compound identification, processing, interpretation, and visualization. LIMS provides a highly specialized, secure, and auditable automation for the experimentation. It covers sample accessioning, preparation, instrumental analysis reporting, and advanced data analysis.

#### Data extraction, compound identification, and curation

Raw data were extracted, peak-identified, and QC-processed with Metabolon’s hardware and software, which are built on a web-service platform utilizing Microsoft’s NET technologies. Metabolites or chemical compounds were identified through a comparison to existing library with the entries of (approximately 3,300) purified authenticated standards and some structurally unnamed biochemicals, which include the retention time/index (RI) within a narrow window, accurate mass to charge ratio (m/z), and chromatographic data (including MS/MS forward and reverse score between the experimental data and authentic standards). The Metabolon’s proprietary visualization and interpretation software was employed to ensure high-quality datasets with accuracy and consistency of peak identification among various samples. Errors from systemic artifacts, mis-assignments, and background noises were corrected or removed for further data quantification and normalization.

#### Metabolite quantification and data normalization

The peaks of chemical compounds were measured using the area under a curve (AUC). A normalization method (termed the “block correction”), in which each compound was corrected in run-day blocks, was introduced for instrumental variations for a multi-day experimental process. In addition, prior to bioinformatical and statistical analysis, data were normalized to total protein content as determined by Bradford assay to account for metabolic differences in each sample.

### PCA of global metabolite profiling

We also used PCA to show global differences between samples. Each principal component is a linear combination of every metabolite, and the principal components are uncorrelated. The number of principal components is equal to the number of observations. PC1 is computed by determining the coefficients of the metabolites that maximizes the variance of the linear combination. PC2 finds the coefficients that maximize the variance with the condition, in which the second component is orthogonal to PC1. PC3 is orthogonal to the first two components. In PCA scatter plots, each data point contains the metabolomic profile of one sample or cell line.

### Hierarchical clustering analysis (HCA) of metabolites

We performed hierarchical clustering, a Euclidean distance-based unsupervised method, where each sample is a vector with all metabolite values, to display large-scale differences among cell lines in a heatmap.

### Pathway enrichment values of global metabolites

In a pair-wise comparison, pathway enrichment can be defined by the number of statistically significantly different metabolites relative to all detected metabolites, compared with the total number of significantly different metabolites relative to all detected metabolites in the study. The pathway enrichment value (*Ev*) is calculated based on the following formula:

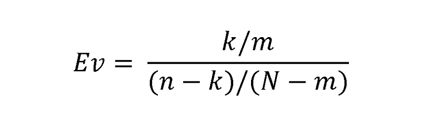

Where: *k* = the number of significant metabolites in the pathway, *m* = the number of metabolites in the pathway, *n* = the total number of significant metabolites, and *N* = the total number of metabolites. An *Ev* greater than one indicates that the pathway holds more significantly changed metabolites relative to all in the study, signifying that the pathway may be a potential area worthy of further investigation.

### Ontological analysis

Enrichr, an intuitive online tool (http://amp.pharm.mssm.edu/Enrichr) (Chen et al., 2013), was used for enrichment ontological analysis of metabolic pathways and their associated gene sets, providing various types of visualization summaries of collective metabolic functions and gene lists (Table S2).

### Standard statistical analysis

For Metabolon datasets, standard statistics was executed in ArrayStudio on log-transformed data or by the statistical programming language R (http://cran.r-project.org/) or JMP data analysis to address significance tests and classifications. One-way ANOVA is used to test whether at least two unknown means are all equal or whether at least one pair of means is different. For the case of two means, ANOVA gives the same result as a two-sided *t*-test with a pooled estimate of the variance. We used two-way ANOVA to compare means among more than two samples and three-way ANOVA to identify biochemicals exhibiting significant interaction and main effects for experimental parameters.

#### Color keys to the pathway explorer map and statistical tables

Green colored values or dots indicate significant differences ( *P* ≤ 0.05) between the groups with metabolite ratios < 1.00; Light Green: narrowly missed statistical cutoff for significance 0.05 < *P* < 0.10, metabolite ratio of < 1.00; Red: indicates significant differences (*P* ≤ 0.05) between the groups with metabolite ratio ≥ 1.00; Light Red: narrowly missed statistical cutoff for significance 0.05 < *P* < 0.10, metabolite ratio of ≥ 1.00; Non-colored text and cell: mean values are not significantly different for that comparison; Blue: indicates significant (*P* ≤ 0.05) ANOVA effects; Light Blue: indicates 0.05 < *P* < 0.10 ANOVA effects.

#### BOX plots

Box plots (e.g., Figure 5C) were used to graphically demonstrate the spread and skewness of numerical data using their quartiles. A boxplot can display at least five elements as follows: the minimum (i.e., the lowest data point in the data set without any outliers), the maximum (the highest data point without any outliers), the sample median (the 50^th^ percentile) as indicated by the horizontal line within the box, the mean value (indicated by a plus sign), the lower quartile (the 25^th^ percentile) showing the median of the lower half of the dataset, and the upper quartile (the 75^th^ percentile) denoting the median of the upper half of the dataset.

## ACKNOWLEDGMENTS

This work was supported by the Intramural Research Program of the National Institutes of Health (NIH) at the National Institute of Neurological Disorders and Stroke (K.G.C., K.P., D.M., K.R.J., B.S.M.) and the National Institute of Dental and Craniofacial Research (P.G.R). We thank Dr. Jason Kinchen at Metabolon Inc., Dr. Hualong Yan at the National Cancer Institute for excellent technical support of this work.

## AUTHOR CONTRIBUTIONS

K.G.C., B.S.M., and P.G.R.: Conceptualization; K.G.C. and B.S.M.: methodology; K.G.C., K.R.J., K.P., D.M., B.S.M: validation and formal analysis; K.G.C., B.S.M.: experimentation and investigation; K.G.C.: writing – original draft; K.G.C., B.S.M., P.G.R.; writing – review and editing; K.G.C.: visualization. All authors approved the final manuscript.

## REFERENCES

Chen EY, Tan CM, Kou Y, Duan Q, Wang Z, Meirelles GV, Clark NR and Ma’ayan A (2013) Enrichr: interactive and collaborative HTML5 gene list enrichment analysis tool. BMC Bioinformatics 14:128.

Chen KG, and Calderone, R. (2016) Adult and cancer stem cells: Perspectives on autophagic fate determinations and molecular intervention, in Targeting Autophagy in Cancer Therapy (Yang JM ed) pp 99-16, Springer International Publishing, Switzerland.

Chen KG, Johnson KR, Park K, Maric D, Yang F, Liu WF, Fann YC, Mallon BS, Robey PG (2024) Resistance to Naïve and Formative Pluripotency Conversion in RSeT Human Embryonic Stem Cells. bioRxiv. doi: 10.1101/2024.02.16.580778

Chen KG, Mallon BS, Hamilton RS, Kozhich OA, Park K, Hoeppner DJ, Robey PG and McKay RD (2012) Non-colony type monolayer culture of human embryonic stem cells. Stem Cell Res 9:237–248.

Chen KG, Mallon BS, Park K, Robey PG, McKay RDG, Gottesman MM and Zheng W (2018) Pluripotent Stem Cell Platforms for Drug Discovery. Trends Mol Med 24:805–820.

Chen KG, Park K and Spence JR (2021) Studying SARS-CoV-2 infectivity and therapeutic responses with complex organoids. Nat Cell Biol 23:822–833.

Cherry AB and Daley GQ (2013) Reprogrammed cells for disease modeling and regenerative medicine. Annu Rev Med 64:277–290.

Collier AJ, Panula SP, Schell JP, Chovanec P, Plaza Reyes A, Petropoulos S, Corcoran AE, Walker R, Douagi I, Lanner F and Rugg-Gunn PJ (2017) Comprehensive Cell Surface Protein Profiling Identifies Specific Markers of Human Naïve and Primed Pluripotent States. Cell Stem Cell 20:874–890 e877.

Cornacchia D, Zhang C, Zimmer B, Chung SY, Fan Y, Soliman MA, Tchieu J, Chambers SM, Shah H, Paull D, Konrad C, Vincendeau M, Noggle SA, Manfredi G, Finley LWS, Cross JR, Betel D and Studer L (2019) Lipid Deprivation Induces a Stable, Naïve-to-Primed Intermediate State of Pluripotency in Human PSCs. Cell Stem Cell 25:120–136 e110.

Diehl FF, Miettinen TP, Elbashir R, Nabel CS, Darnell AM, Do BT, Manalis SR, Lewis CA and Vander Heiden MG (2022) Nucleotide imbalance decouples cell growth from cell proliferation. Nat Cell Biol 24:1252–1264.

Dong C, Fischer LA and Theunissen TW (2019) Recent insights into the naïve state of human pluripotency and its applications. Exp Cell Res 385:111645.

Fendt SM, Bell EL, Keibler MA, Olenchock BA, Mayers JR, Wasylenko TM, Vokes NI, Guarente L, Vander Heiden MG and Stephanopoulos G (2013) Reductive glutamine metabolism is a function of the alpha-ketoglutarate to citrate ratio in cells. Nat Commun 4:2236.

Forristal CE, Wright KL, Hanley NA, Oreffo RO and Houghton FD (2010) Hypoxia inducible factors regulate pluripotency and proliferation in human embryonic stem cells cultured at reduced oxygen tensions. Reproduction 139:85–97.

Gafni O, Weinberger L, Mansour AA, Manor YS, Chomsky E, Ben-Yosef D, Kalma Y, Viukov S, Maza I, Zviran A, Rais Y, Shipony Z, Mukamel Z, Krupalnik V, Zerbib M, Geula S, Caspi I, Schneir D, Shwartz T, Gilad S, Amann-Zalcenstein D, Benjamin S, Amit I, Tanay A, Massarwa R, Novershtern N and Hanna JH (2013) Derivation of novel human ground state naïve pluripotent stem cells. Nature 504:282–286.

Johnson KR, Mallon BS, Fann YC and Chen KG (2021) Multivariate meta-analysis reveals global transcriptomic signatures underlying distinct human naïve-like pluripotent states. PLoS One 16:e0251461.

Kilens S, Meistermann D, Moreno D, Chariau C, Gaignerie A, Reignier A, Lelievre Y, Casanova M, Vallot C, Nedellec S, Flippe L, Firmin J, Song J, Charpentier E, Lammers J, Donnart A, Marec N, Deb W, Bihouee A, Le Caignec C, Pecqueur C, Redon R, Barriere P, Bourdon J, Pasque V, Soumillon M, Mikkelsen TS, Rougeulle C, Freour T, David L and Milieu Interieur C (2018) Parallel derivation of isogenic human primed and naïve induced pluripotent stem cells. Nat Commun 9:360.

Lees JG, Cliff TS, Gammilonghi A, Ryall JG, Dalton S, Gardner DK and Harvey AJ (2019) Oxygen Regulates Human Pluripotent Stem Cell Metabolic Flux. Stem Cells Int 2019:8195614.

Liu X, Nefzger CM, Rossello FJ, Chen J, Knaupp AS, Firas J, Ford E, Pflueger J, Paynter JM, Chy HS, O’Brien CM, Huang C, Mishra K, Hodgson-Garms M, Jansz N, Williams SM, Blewitt ME, Nilsson SK, Schittenhelm RB, Laslett AL, Lister R and Polo JM (2017) Comprehensive characterization of distinct states of human naïve pluripotency generated by reprogramming. Nat Methods 14:1055–1062.

Mallon BS, Chenoweth JG, Johnson KR, Hamilton RS, Tesar PJ, Yavatkar AS, Tyson LJ, Park K, Chen KG, Fann YC and McKay RD (2013) StemCellDB: the human pluripotent stem cell database at the National Institutes of Health. Stem Cell Res 10:57–66.

Mathews CK (2015) Deoxyribonucleotide metabolism, mutagenesis and cancer. Nat Rev Cancer 15:528–539.

McCoin CS, Knotts TA and Adams SH (2015) Acylcarnitines--old actors auditioning for new roles in metabolic physiology. Nat Rev Endocrinol 11:617–625.

Metallo CM, Gameiro PA, Bell EL, Mattaini KR, Yang J, Hiller K, Jewell CM, Johnson ZR, Irvine DJ, Guarente L, Kelleher JK, Vander Heiden MG, Iliopoulos O and Stephanopoulos G (2011) Reductive glutamine metabolism by IDH1 mediates lipogenesis under hypoxia. Nature 481:380–384.

Rubinsztein DC, Marino G and Kroemer G (2011) Autophagy and aging. Cell 146:682–695. Smith A (2017) Formative pluripotency: the executive phase in a developmental continuum. Development 144:365-373.

Somasundaram L, Levy S, Hussein AM, Ehnes DD, Mathieu J and Ruohola-Baker H (2020) Epigenetic metabolites license stem cell states. Curr Top Dev Biol 138:209–240.

Spyrou J, Gardner DK and Harvey AJ (2019) Metabolomic and Transcriptional Analyses Reveal Atmospheric Oxygen During Human Induced Pluripotent Stem Cell Generation Impairs Metabolic Reprogramming. Stem Cells 37:1042–1056.

Szczerbinska I, Gonzales KAU, Cukuroglu E, Ramli MNB, Lee BPG, Tan CP, Wong CK, Rancati GI, Liang H, Goke J, Ng HH and Chan YS (2019) A Chemically Defined Feeder-free System for the Establishment and Maintenance of the Human Naïve Pluripotent State. Stem Cell Reports 13:612–626.

Takashima Y, Guo G, Loos R, Nichols J, Ficz G, Krueger F, Oxley D, Santos F, Clarke J, Mansfield W, Reik W, Bertone P and Smith A (2014) Resetting transcription factor control circuitry toward ground-state pluripotency in human. Cell 158:1254–1269.

Theunissen TW, Powell BE, Wang H, Mitalipova M, Faddah DA, Reddy J, Fan ZP, Maetzel D, Ganz K, Shi L, Lungjangwa T, Imsoonthornruksa S, Stelzer Y, Rangarajan S, D’Alessio A, Zhang J, Gao Q, Dawlaty MM, Young RA, Gray NS and Jaenisch R (2014) Systematic identification of culture conditions for induction and maintenance of naïve human pluripotency. Cell Stem Cell 15:471–487.

Tsogtbaatar E, Landin C, Minter-Dykhouse K and Folmes CDL (2020) Energy Metabolism Regulates Stem Cell Pluripotency. Front Cell Dev Biol 8:87.

Weinberger L, Ayyash M, Novershtern N and Hanna JH (2016) Dynamic stem cell states: naïve to primed pluripotency in rodents and humans. Nat Rev Mol Cell Biol 17:155–169.

Wu J and Izpisua Belmonte JC (2015) Metabolic exit from naïve pluripotency. Nat Cell Biol 17:1519–1521.

Zhang J, Zhao J, Dahan P, Lu V, Zhang C, Li H and Teitell MA (2018) Metabolism in Pluripotent Stem Cells and Early Mammalian Development. Cell Metab 27:332–338.

Zhou J, Hu J, Wang Y and Gao S (2023) Induction and application of human naïve pluripotency. Cell Rep 42:112379.

Zhou W, Choi M, Margineantu D, Margaretha L, Hesson J, Cavanaugh C, Blau CA, Horwitz MS, Hockenbery D, Ware C and Ruohola-Baker H (2012) HIF1alpha induced switch from bivalent to exclusively glycolytic metabolism during ESC-to-EpiSC/hESC transition. EMBO J 31:2103–2116.

Zimmerlin L, Park TS and Zambidis ET (2017) Capturing Human Naïve Pluripotency in the Embryo and in the Dish. Stem Cells Dev 26:1141–1161.

