## Supplementary figures and images for "Metabolic Quadrivalency in RSeT Human Embryonic Stem Cells"

### Figure S1

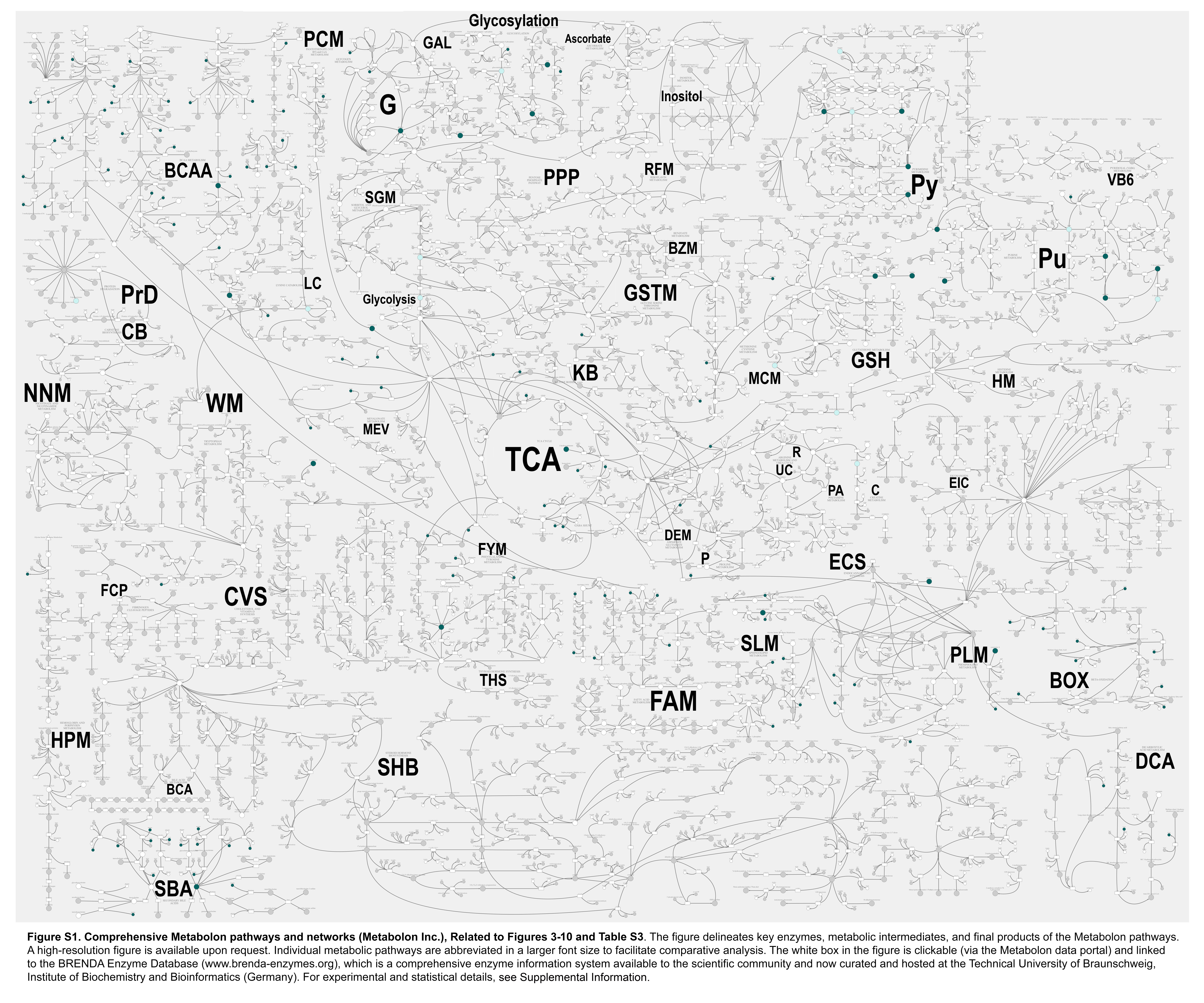

### Figure S2

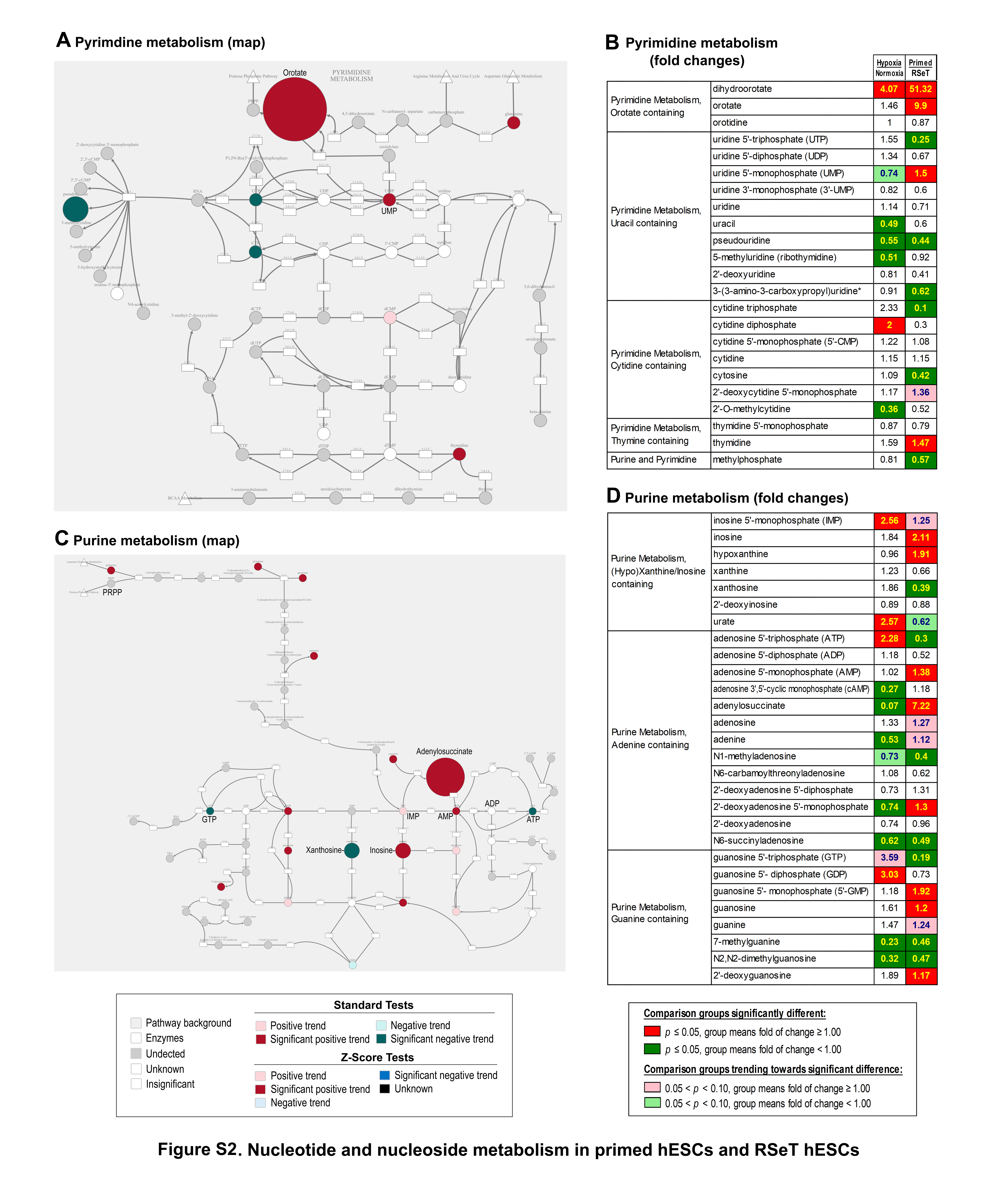
