## Supplementary material for "Metabolic Quadrivalency in RSeT Human Embryonic Stem Cells": Table S1

**Table S1.** **Gene expression signatures underlying metabolism in human pluripotent stem cells**

| **Gene Symbols** | **Description**  **(Based on GeneCards and**  **Entrez of NCBI)** | **Gene expression**  **probes** | **RSeT**  **vs primed**  **(n = 6, Fold changes)** | **RN vs PN**  **(n = 3, fold changes)** | **RH vs PH**  **(n = 3,**  **fold changes)** |
| --- | --- | --- | --- | --- | --- |
| **Metabolism gene expression** (n = 93, ** indicates *P* < 0.05) | | | | | |
| *AADAT* | Aminoadipate Aminotransferase | A_23_P253484 | 0.61 | 0.59 | 0.63 |
| *AASS* | Aminoadipate-Semialdehyde Synthase | A_23_P8754 | 0.63 | 0.83 | 0.48 |
| *ACACA* | Acetyl-CoA Carboxylase Alpha | A_32_P215318 | **2.86*** | **2.95*** | **2.76*** |
| *ACLY* | ATP Citrate Lyase | A_23_P66787 | **1.47*** | 1.31 | 1.65 |
| *ACO2* | Aconitase 2 | A_23_P103149 | **1.64*** | **1.51*** | **1.79*** |
| *AGMAT* | Agmatinase | A_23_P103720 | 0.90 | 1.02 | 0.80 |
| *AHCY* | Adenosylhomocysteinase | A_23_P17575 | **1.17*** | **1.21*** | 1.12 |
| *AHCYL2* | Adenosylhomocysteinase Like 2 | A_24_P72518 | **1.39*** | 1.34 | 1.45 |
| *ALDH1A1* | Aldehyde Dehydrogenase 1  Family Member A1 | A_23_P83098 | 0.51 | **0.26*** | 1.00 |
| *ALDH1B1* | Aldehyde Dehydrogenase 1  Family Member B1 | A_23_P135294 | **0.76*** | **0.63*** | 0.91 |
| *ALOX15B* | Arachidonate 15-Lipoxygenase Type B | A_23_P60627 | 1.86 | 1.29 | 2.70 |
| *ALOX5* | Arachidonate 5-Lipoxygenase | A_23_P104464 | 1.22 | 2.08 | 0.72 |
| *AOX1* | Aldehyde Oxidase 1 | A_23_P154037 | **0.69*** | 0.64 | 0.75 |
| *ASL* | Argininosuccinate Lyase | A_23_P26223 | 0.95 | 0.80 | 1.11 |
| *ASS1* | Argininosuccinate Synthase 1 | A_23_P31921 | 1.03 | 0.88 | 1.19 |
| *ATP5E* | ATP5F1E, ATP Synthase F1  Subunit Epsilon | A_23_P252322 | **0.62*** | **0.66*** | **0.57*** |
| *ATP5F1* | ATP5PB, ATP Synthase Peripheral Stalk-Membrane Subunit B | A_23_P46275 | **1.55*** | 1.93 | **1.24*** |
| *ATP6* | MT-ATP6, Mitochondrially Encoded ATP Synthase Membrane Subunit 6 | A_23_P337726 | 1.04 | 0.99 | 1.10 |
| *BCAT1* | Branched Chain Amino Acid Transaminase 1 | A_24_P935986 | **0.62*** | **0.65*** | 0.60 |
| *BDH2* | 3-Hydroxybutyrate Dehydrogenase 2 | A_23_P92490 | 0.93 | **0.94*** | 0.92 |
| *BHMT* | Betaine--Homocysteine S-Methyltransferase | A_23_P81581 | **2.13*** | 2.71 | 1.67 |
| *COX6B2* | Cytochrome C Oxidase Subunit 6B2 | A_24_P267523 | 1.95 | 2.06 | 1.85 |
| *CPS1* | Carbamoyl-Phosphate Synthase 1 | A_23_P5300 | **1.98*** | 1.94 | 2.03 |
| *CS* | Citrate Synthase | A_23_P47818 | **1.55*** | **1.25*** | **1.91*** |
| *CYP4F2* | Cytochrome P450 Family 4 Subfamily F Member 2 | A_24_P168494 | 0.83 | 0.83 | 0.83 |
| *DBT* | Dihydrolipoamide Branched Chain Transacylase E2 | A_24_P28524 | **1.35*** | 1.38 | **1.31*** |
| *DHH* | Desert Hedgehog Signaling Molecule | A_23_P162314 | 0.82 | 0.77 | 0.88 |
| *DLAT* | Dihydrolipoamide S-Acetyltransferase | A_24_P372672 | **1.42*** | 1.28 | **1.58*** |
| *DLD* | Dihydrolipoamide Dehydrogenase | A_23_P111835 | **1.28*** | 1.15 | **1.43*** |
| *ENO1* | Enolase 1 | A_24_P105501 | 0.98 | 0.87 | 1.11 |
| *ENO1-AS1* | ENO1 Antisense RNA 1 | A_32_P145447 | 0.90 | 0.84 | 0.97 |
| *ENO3* | Enolase 3 | A_23_P130149 | **1.19*** | 1.26 | 1.13 |
| *FA2H* | Fatty Acid 2-Hydroxylase | A_23_P49448 | **6.12*** | **6.07*** | **6.18*** |
| *FAAH* | Fatty Acid Amide Hydrolase | A_23_P103226 | **0.31*** | **0.25*** | **0.38*** |
| *FASN* | Fatty Acid Synthase | A_23_P44132 | **2.48*** | 1.85 | 3.31 |
| *GAD1* | Glutamate Decarboxylase 1 | A_23_P209578 | 0.54 | 0.57 | 0.50 |
| *GLS* | Glutaminase | A_23_P308800 | **0.50*** | 0.67 | 0.37 |
| *GLUD1* | Glutamate Dehydrogenase 1 | A_23_P138665 | **1.66*** | 1.64 | 1.69 |
| *GLUD2* | Glutamate Dehydrogenase 2 | A_23_P45361 | 1.62 | 1.30 | 2.01 |
| *GOT1* | Glutamic-Oxaloacetic Transaminase 1 | A_23_P63825 | **1.31*** | **1.24*** | 1.38 |
| *GOT2* | Glutamic-Oxaloacetic Transaminase 2 | A_23_P106575 | 0.96 | 0.83 | 1.12 |
| *GPD1* | Glycerol-3-Phosphate Dehydrogenase 1 | A_23_P413721 | 0.97 | 0.91 | 1.03 |
| *GPD1L* | Glycerol-3-Phosphate Dehydrogenase 1 Like | A_23_P318284 | **0.42*** | 0.48 | **0.38*** |
| *GPT* | Glutamic--Pyruvic Transaminase | A_23_P146339 | 1.07 | 1.23 | 0.93 |
| *GYS1* | Glycogen Synthase 1 | A_23_P208698 | 1.27 | 1.03 | 1.56 |
| *HAGHL* | Hydroxyacylglutathione Hydrolase Like | A_24_P356373 | 0.86 | 0.74 | 1.00 |
| *HMGCS1* | 3-Hydroxy-3-Methylglutaryl-CoA Synthase 1 | A_23_P133263 | **1.50*** | **1.51*** | **1.49*** |
| *IDH1* | Isocitrate Dehydrogenase (NADP^+^) 1 | A_32_P45009 | 1.13 | 1.04 | 1.23 |
| *IDH1-AS1* | IDH1 Antisense RNA 1 | A_32_P16931 | 1.42 | 1.79 | 1.13 |
| *IDH3A* | Isocitrate Dehydrogenase [NAD^+^] 3 Catalytic Subunit Alpha | A_23_P140668 | 1.65 | 1.42 | 1.91 |
| *IDO1* | Indoleamine 2,3-Dioxygenase 1 | A_23_P112026 | **0.63*** | 0.69 | **0.57*** |
| *IL4I1* | Interleukin 4 Induced 1 | A_23_P502520 | 1.26 | 1.36 | 1.17 |
| *IMPA2* | Inositol Monophosphatase 2 | A_23_P50081 | **0.61*** | **0.60*** | **0.62*** |
| ***LDHC**** | Lactate Dehydrogenase C  **Transcriptionally regulated in mESCs** | A_23_P53039 | 1.17 | 1.34 | 1.03 |
| *MAOA* | Monoamine Oxidase A | A_23_P83857 | **0.56*** | 0.49 | 0.64 |
| *MAOB* | Monoamine Oxidase B | A_23_P85008 | 1.05 | 1.09 | 1.00 |
| *MAT1A* | Methionine Adenosyltransferase 1A | A_23_P23996 | **0.26*** | 0.25 | 0.27 |
| *NAT8L* | N-Acetyltransferase 8 Like | A_24_P91991 | **0.50*** | **0.61*** | 0.41 |
| *NMRK1* | Nicotinamide Riboside Kinase 1 | A_23_P32036 | **0.79*** | 0.78 | **0.80*** |
| *NOS1AP* | Nitric Oxide Synthase 1 Adaptor Protein | A_23_P74309 | **2.19*** | 1.93 | 2.48 |
| *NOS2* | Nitric Oxide Synthase 2 | A_23_P502464 | 1.16 | 2.09 | 0.64 |
| *NT5C2* | 5'-Nucleotidase, Cytosolic II | A_23_P97906 | **0.47*** | **0.54*** | **0.40*** |
| *ODC1* | Ornithine Decarboxylase 1 | A_23_P165840 | 1.59 | 1.44 | 1.74 |
| *OGDH* | Oxoglutarate Dehydrogenase | A_23_P123133 | 1.66 | 1.64 | 1.68 |
| *PC* | Pyruvate Carboxylase | A_23_P161647 | 1.00 | 1.07 | 0.94 |
| *PDHA1* | Pyruvate Dehydrogenase E1 Subunit Alpha 1 | A_23_P251095 | **1.57*** | **1.51*** | **1.63*** |
| *PDHB* | Pyruvate Dehydrogenase E1 Subunit Beta | A_23_P20932 | 0.85 | **0.75*** | 0.97 |
| *PDK1* | Pyruvate Dehydrogenase Kinase 1 | A_23_P10614 | **1.59*** | 1.69 | **1.49*** |
| *PDK2* | Pyruvate Dehydrogenase Kinase 2 | A_23_P207517 | 0.94 | **0.72*** | 1.23 |
| *PDK3* | Pyruvate Dehydrogenase Kinase 3 | A_32_P214503 | 1.04 | **1.13*** | 0.95 |
| ***PDXDC1*** | Pyridoxal Dependent Decarboxylase Domain Containing 1  **Transcriptionally regulated in mESCs, high in Tesar data** | A_23_P391583 | **0.63*** | **0.53*** | **0.74*** |
| *PFKL* | Phosphofructokinase, Liver Type | A_23_P29079 | 2.18 | 1.94 | 2.44 |
| *PGK1* | Phosphoglycerate Kinase 1 | A_24_P254532 | 1.58 | 1.87 | 1.33 |
| ***PLA2G10*** | Phospholipase A2 Group X;  **Transcriptionally regulated in mESCs, high in Tesar data** | A_23_P88767 | **2.87*** | 2.73 | 3.02 |
| *PPARGC1B* | PPARG Coactivator 1 Beta  (Known as PGC-1-Beta) | A_24_P560519 | **2.23*** | **2.05*** | 2.44 |
| *PRKACA* | Protein Kinase cAMP-Activated Catalytic Subunit Alpha | A_24_P399630 | 1.58 | 1.27 | 1.98 |
| *PRODH* | Proline Dehydrogenase 1 | A_23_P68786 | **5.65*** | **3.38*** | **9.45*** |
| *PYCR1* | Pyrroline-5-Carboxylate Reductase 1 | A_23_P130194 | **0.49*** | 0.48 | **0.50*** |
| *PYCR2* | Pyrroline-5-Carboxylate Reductase 2 | A_24_P165259 | **0.48*** | 0.63 | 0.37 |
| ***PYGL*** | Glycogen Phosphorylase L  **Transcriptionally regulated with naïve protocols,** low in naïve mESCs | A_23_P48676 | 0.65 | 0.67 | 0.64 |
| *PYGM* | Glycogen Phosphorylase, Muscle Associated | A_23_P52657 | 1.24 | 1.32 | 1.17 |
| *RRM2B* | Ribonucleotide Reductase Regulatory TP53 Inducible Subunit M2B | A_23_P20225 | **0.56*** | **0.76*** | **0.42*** |
| *SC5D* | Sterol-C5-Desaturase | A_23_P98446 | 0.83 | 0.73 | 0.94 |
| *SCO2* | Synthesis of Cytochrome C Oxidase 2 | A_23_P132388 | **0.77*** | 0.80 | 0.73 |
| *SDHB* | Succinate Dehydrogenase Complex Iron Sulfur Subunit B | A_23_P149649 | 1.09 | 0.99 | 1.20 |
| *SDS* | Serine Dehydratase | A_24_P304439 | **0.52*** | 0.45 | 0.60 |
| ***SLC16A2*** | Monocarboxylate Transporter;  **Transcriptionally regulated in mESCs, high in Tesar data** | A_23_P137097 | **3.04*** | **2.55*** | **3.62*** |
| *SLC29A1* | Solute Carrier Family 29 Member 1 (Augustine Blood Group) | A_23_P133694 | **0.76*** | **0.67*** | 0.86 |
| *SMS* | Spermine Synthase | A_32_P50452 | 1.02 | 1.08 | 0.96 |
| *SQLE* | Squalene Epoxidase | A_23_P146284 | **1.41*** | **1.35*** | 1.48 |
| *TDH* | L-Threonine Dehydrogenase (Pseudogene) | A_23_P337778 | **4.47*** | **3.59*** | **5.56*** |
| *TK2* | Thymidine Kinase 2 | A_23_P351679 | 1.15 | 1.36 | 0.97 |
| *TYMP* | Thymidine Phosphorylase | A_23_P91802 | **0.56*** | 0.63 | 0.69 |

**FOOTNOTES**

**^*^**Transcriptionally regulated genes are indicated in bold gene symbols, with up-regrated gene in red and down-regulated gene in blue colors.

****** Indicates *P* values < 0.05 with two-tailed Student *t*-test, for fold changes of gene expression in microarray.

Abbreviations: mESCs, naive mouse embryonic stem cells; PN and PH, primed hESCs under normoxic and hypoxic growth conditions respectively; RN and RH, RSeT hESCs under normoxic and hypoxic growth conditions respectively; Tesar data: data published by Tesar et al., Nature 2007 (PMID: 17597760).
